# Spatial immune architecture and tumor lineage programs jointly shape clinical outcomes in advanced pancreatic ductal adenocarcinoma

**DOI:** 10.64898/2026.06.28.734525

**Authors:** Hay Mar Oo, Tauangtham Anekpuritanang, Napat Angkathunyakul, Ufuk Degirmenci, Ananya Pongpaibul, Siwakorn Punyawatthananukool, Krittiya Korphaisarn, Somponnat Sampattavanich

**Affiliations:** Siriraj Center of Research Excellence for Systems Pharmacology, Department of Pharmacology, Faculty of Medicine Siriraj Hospital, Mahidol University, Bangkok, 10700, Thailand; Department of Pathology, Department of Medicine, Faculty of Medicine Siriraj Hospital, Mahidol University, Bangkok, 10700, Thailand; Spatial Biology Department, Next Level Genomics, 138672, Singapore; Department of Pharmacology, National University of Singapore, 119077, Singapore; Division of Medical Oncology, Department of Medicine, Faculty of Medicine Siriraj Hospital, Mahidol University, Bangkok, 10700, Thailand

**Keywords:** Pancreatic ductal adenocarcinoma, spatial biology, tumor lineage states, spatial immune organization, epithelial plasticity, tumor-associated macrophages, prognostic biomarkers

## Abstract

Pancreatic ductal adenocarcinoma (PDAC) exhibits extensive molecular and microenvironmental heterogeneity, yet how tumor lineage states interact with spatial immune organization in advanced disease remains poorly understood. Here, we performed multiplexed spatial proteomic profiling using tissue cyclic immunofluorescence (t-CyCIF) in 27 patients with advanced PDAC and integrated these analyses with spatial transcriptomic profiling of representative tumors. Tumors were classified into Classical, Hybrid, Basal, and Null epithelial states based on GATA6 and CK5 expression, revealing distinct immune architectures associated with clinical outcome. Classical and Hybrid tumors displayed immune-inflamed microenvironments enriched for lymphocytes, whereas Basal and Null tumors exhibited immune-excluded, macrophage-dominated landscapes characterized by increased M2 macrophages.

Spatial transcriptomic analysis further revealed that Hybrid tumors were not homogeneous intermediate states but instead contained spatially segregated Hybrid_Classical and Hybrid_Basal regions with distinct transcriptional programs, immune niches, and cell–cell communication networks. Hybrid_Basal regions were associated with increased M2 macrophage enrichment and preferential activation of macrophage-derived SPP1–CD44 signaling, implicating localized immune–epithelial interactions in epithelial plasticity and lineage-state transitions.

To quantify spatial immune organization, we developed a spatial immune score that captures the relative positioning of CD8⁺ cytotoxic T cells with respect to CD4⁺ helper T cells and CD163⁺ M2 macrophages. Higher scores were associated with worse survival and provided stronger prognostic information than conventional immune cell abundance metrics. Integration of the spatial immune score with GATA6 expression achieved superior prognostic discrimination (AUC = 0.822) compared with either feature alone. Together, these findings demonstrate that tumor lineage state and spatial immune organization represent complementary dimensions of PDAC biology and highlight spatial tumor–immune interactions as determinants of clinical outcome in advanced pancreatic cancer.

**Summary:** Pancreatic ductal adenocarcinoma (PDAC) exhibits marked molecular and microenvironmental heterogeneity, yet how tumor lineage states interact with the spatial immune microenvironment in advanced disease remains poorly understood. Here, the authors apply multiplexed spatial proteomics and spatial transcriptomics to advanced PDAC and show that epithelial lineage states defined by GATA6 and CK5 are associated with distinct immune architectures and macrophage-enriched signaling niches. Hybrid tumors contain spatially segregated epithelial states with differential immune engagement and SPP1–CD44 signaling. The authors further identify a spatial immune score based on the relative positioning of CD8⁺ T cells, CD4⁺ T cells, and M2 macrophages that predicts patient survival. Integration of spatial immune organization with tumor lineage information improves prognostic stratification, highlighting the clinical relevance of spatial tumor–immune interactions in advanced PDAC.

## Introduction

Pancreatic ductal adenocarcinoma (PDAC) remains one of the most lethal malignancies worldwide, with five-year survival rates below 10% for patients diagnosed with advanced disease^1^. Despite extensive genomic and transcriptomic characterization, clinical outcomes remain highly heterogeneous and are poorly predicted by tumor-intrinsic features alone^2^. Increasing evidence indicates that tumor progression and therapeutic resistance are strongly influenced by the tumor microenvironment (TME), particularly immune and stromal interactions^3, 4^. However, most studies have analyzed tumor molecular subtypes and immune composition independently, often relying on bulk transcriptomic approaches that lack spatial resolution. Consequently, how tumor cell identity interacts with the spatial organization of immune cells within the tumor microenvironment and how this relationship influences patient outcomes remains poorly understood. Addressing this gap is particularly critical for advanced PDAC, where most patients are diagnosed at unresectable stages^3^, yet spatial profiling studies have largely focused on surgically resected tumors^5, 6^.

Molecular subtypes, particularly classical and basal-like, are strongly associated with clinical outcomes and chemotherapy response in PDAC. The classical subtype, characterized by high GATA6 expression, is associated with longer survival and increased sensitivity to FOLFIRINOX, whereas the basal-like subtype, marked by CK5 expression, is linked to poorer prognosis; evaluated in the COMPASS study (NCT02750657), which assessed the predictive value of these subtypes for survival and treatment response to first-line modified FOLFIRINOX and gemcitabine plus nab-paclitaxel in advanced PDAC^2, 7, 8, 9^. However, accumulating evidence indicates substantial subtype plasticity, with tumors transitioning between classical and basal-like states and giving rise to intermediate or hybrid phenotypes that contribute to heterogeneity in treatment response^10, 11, 12, 13^. Notably, chemotherapy itself can promote a shift from classical to basal-like transcriptional states, a transition associated with increased chemoresistance and reduced patient survival^14, 15, 16, 17^. Together, these findings suggest the intra-tumoral subtype heterogeneity behavior of PDAC necessitates spatially resolved approaches for molecular classification.

The tumor microenvironment (TME) also plays a critical role in tumor progression and therapy resistance in PDAC^18^. Recent studies have identified distinct immune contexts associated with PDAC molecular states, including reactive and deserted tumor microenvironments linked to basal and classical subtypes, respectively^19^. Moreover, tumor-extrinsic signals can actively shape tumor cell states. For example, macrophage-derived cytokines such as TGF-β and TNF-α have been shown to induce basal-like transcriptional programs by regulating tumor-intrinsic gene expression networks^13, 20, 21^. Despite these advances, a fundamental gap persists in our understanding of how tumor molecular subtypes and immune landscapes are spatially organized and functionally interrelated at single-cell resolution. As a result, how heterogeneous tumor cell states are spatially associated with immune architecture within individual tumors, and whether these spatial relationships contribute to clinical outcomes, remain poorly understood.

GATA6 and CK5 have been established as robust lineage markers of classical and basal-like PDAC states, respectively, with GATA6⁺ tumors generally exhibiting more differentiated epithelial phenotypes, enriched immune infiltration, and improved clinical outcomes, whereas CK5⁺ tumors display basal-like features associated with aggressive behavior and poorer prognosis^22^. These markers have also demonstrated prognostic value when assessed by immunohistochemistry in patients receiving neoadjuvant chemotherapy^23^. Nonetheless, conventional single-marker immunohistochemistry lacks the spatial resolution required to capture the organization of tumor and immune cells within the tissue microenvironment. Consequently, it remains difficult to determine how tumor lineage states interact with immune cell positioning and cell–cell interactions to shape functional immune niches and influence patient prognosis.

Here, we apply highly multiplexed tissue cyclic immunofluorescence (t-CyCIF) to profile the spatial immune landscape of advanced-stage (III/IV) PDAC at single-cell resolution. By integrating GATA6- and CK5-based tumor lineage classification with microenvironmental profiling, we map tumor cell states, immune composition, and their spatial organization within the tumor microenvironment. We identify a spatial immune positioning axis associated with patient outcomes and show that integrating this spatial metric with tumor lineage state enhances prognostic discrimination beyond molecular subtyping alone. These findings establish tumor lineage and spatial immune architecture as complementary determinants of disease behavior and provide a framework for spatially informed prognostic stratification in advanced PDAC.

## Results

### Multiplex spatial profiling identifies PDAC molecular subtypes in advanced disease

To characterize tumor cell states and their associated immune architecture in advanced pancreatic ductal adenocarcinoma (PDAC), we performed multiplex spatial profiling using tissue cyclic immunofluorescence (t-CyCIF) on tumor specimens from 27 patients with stage III–IV disease (Fig. 1a). The cohort included both primary pancreatic tumors (n = 16) and metastatic lesions from liver (n = 7), peritoneum (n = 2), omentum (n = 1), and stomach (n = 1), enabling analysis across clinically relevant sites of advanced disease. Tumor epithelial cells were identified as pan-cytokeratin–positive (PK⁺) cells and classified based on the expression of the lineage markers GATA6 and CK5. Using these markers, tumors were segregated into four molecular subtypes: classical (GATA6⁺CK5⁻, n = 8), basal (GATA6⁻CK5⁺, n = 5), hybrid (GATA6⁺CK5⁺, n = 7), and null (GATA6⁻CK5⁻, n = 7) (Fig. 1b, d). Subtype classification was assigned according to cohort-derived mean expression thresholds, and subtype-specific scores derived from PK⁺ tumor cells clearly separated these groups (Fig. 1c). Consistent with this classification, the relative proportions of GATA6⁺, CK5⁺, and double-positive tumor cell populations differed significantly across subtypes (p < 0.05; Supplementary Fig. 1a), supporting the robustness of molecular subtyping in our cohort. We also observed substantial heterogeneity in the proportions of GATA6⁺, CK5⁺, and double-positive tumor cells both across patients and within individual tumors (Fig. 1e), highlighting pronounced intra- and inter-tumor lineage diversity.

**Fig. 1.**
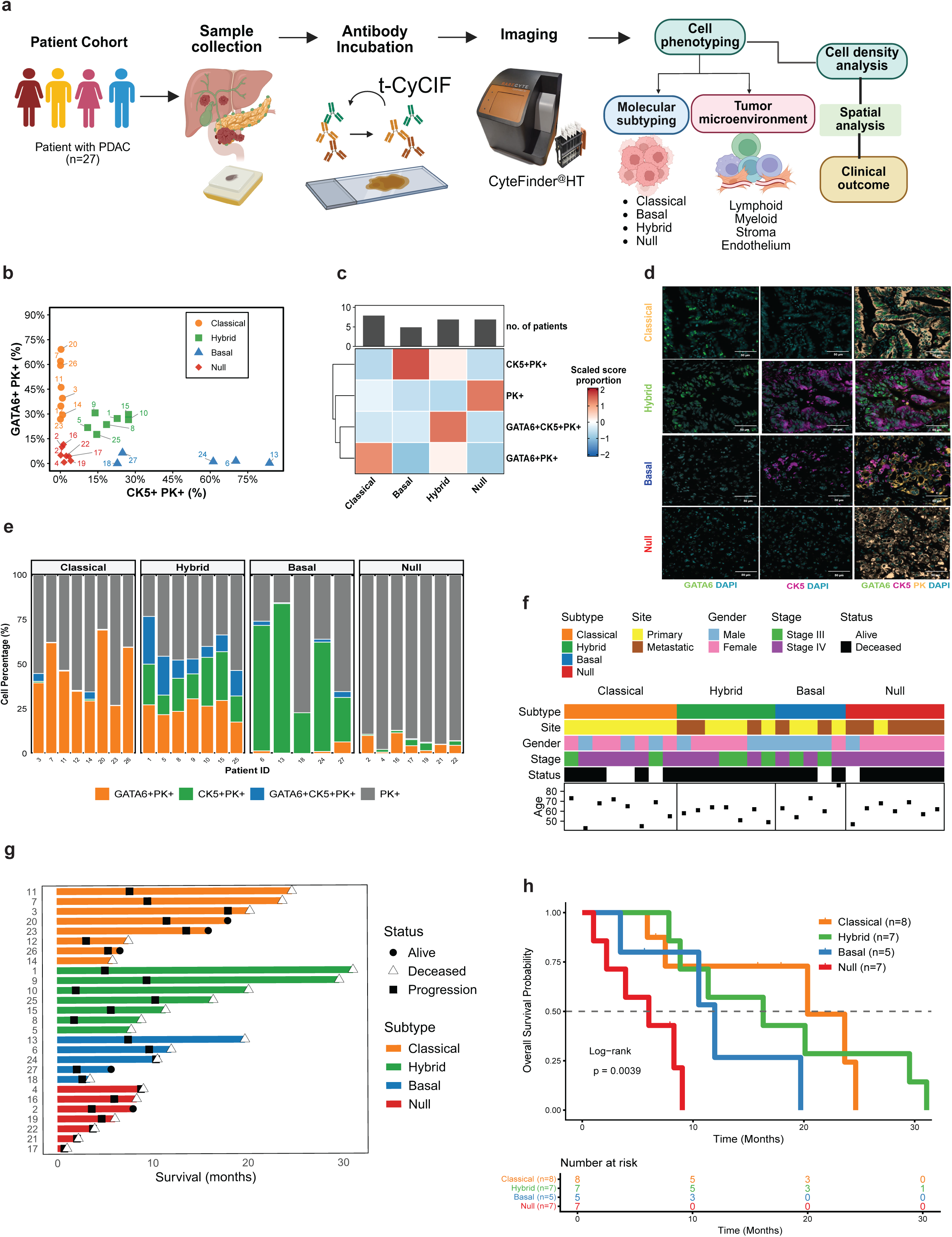
Multiplex spatial profiling identifies PDAC molecular subtypes in advanced disease. **a** Overall experimental workflow. Sample collection, t-CyCIF method with imaging and further data analysis. This figure was created in https://BioRender.com. **b** Scatter plot showing GATA6⁺PK⁺ versus CK5⁺PK⁺ cell proportions to distinguish the molecular subtypes (n=27). **c** Heatmap of mean expression Z-scores of nuclear GATA6 and cytoplasmic CK5 proportions across 27 PDAC patients (n = 27), used for GATA6/CK5-based tumor subtype classification. **d** Representative multiplex immunofluorescence images showing nuclear GATA6 (green), cytoplasmic CK5 (magenta), and pan-cytokeratin (PK, yellow) across four PDAC subtypes. Scale bars, 50 μm. **e** Stacked bar charts showing molecular marker proportions per patient such as Classical (GATA6^+^PK^+^), Hybrid (GATA6^+^CK5^+^PK^+^), Basal (GATA6^-^CK5^+^PK^+^), Null (GATA6^-^CK5^-^PK^+^). **f** Clinical information for patient samples including age, status, tumor stage (III, IV) at diagnosis, gender, site of specimen archived (primary or metastasis), subtypes information (Classical, Hybrid, Basal, Null). **g** Swimmer plot showing individual patient survival durations (Alive, deceased, progression) by molecular subtypes. **h** Kaplan–Meier overall survival curves by subtypes and log-rank test was also used for survival analysis. *P* values < 0.05 were considered statistically significant. All analysis was done in all patients (n=27).

Individual patient trajectories, including time-to-event and censoring status (alive, deceased, or progression), are shown in Fig. 1g, providing the underlying event data used to construct the Kaplan–Meier survival curves in Fig. 1h. Kaplan–Meier analysis demonstrated significant differences in overall survival among the four molecular subtypes (log-rank P = 0.0039; Fig. 1h), with classical and hybrid tumors associated with longer median overall survival (20.33 and 16.32 months, respectively) compared to basal and null tumors (11.95 and 6.01 months, respectively). In contrast, clinical variables including age, gender, tumor site, and disease stage were not significantly associated with overall survival (Fig. 1f; Supplementary Fig. 2a; Table 1). Consistently, univariable Cox proportional hazards analysis identified molecular subtype as a strong predictor of clinical outcome, with the null subtype exhibiting the highest risk relative to the classical subtype (hazard ratio (HR) = 6.38, 95% confidence interval (CI) 1.88–21.7, p = 0.003; Supplementary Fig. 2b).

**Table 1.** Clinicopathological characteristics.

| **Characteristic** | **n = 27** |
| --- | --- |
| Mean age at diagnosis (IQR) | 61 (43, 86) |
| **Age group** |  |
| <60 | 12 (44%) |
| ≥60 | 15 (56%) |
| **Sex** |  |
| Male | 11 (41%) |
| Female | 16 (59%) |
| **ECOG performance status** |  |
| 0–1 | 26 (96%) |
| ≥2 | 1 (3.7%) |
| **Molecular subtypes** |  |
| Classical | 8 (30%) |
| Hybrid | 7 (26%) |
| Basal | 5 (19%) |
| Null | 7 (26%) |
| **Stage at diagnosis** |  |
| Locally advanced (stage III) | 7 (26%) |
| Metastatic (stage IV) | 20 (74%) |
| **Specimen site (type of specimen)** |  |
| Primary site | 16 (59%) |
| Metastatic site | 11 (41%) |
| **CA19-9 category** |  |
| Normal | 8 (30%) |
| Elevated <59 U/mL | 13 (48%) |
| Elevated ≥59 U/mL | 6 (22%) |
| **First-line regimen** |  |
| FOLFIRINOX | 15 (56%) |
| Gemcitabine/Capecitabine | 12 (44%) |

Data are n (%) or mean (range). ECOG, Eastern Cooperative Oncology Group. All patients presented with locally advanced or metastatic disease. Specimen site denotes the tissue source used for t-CyCIF staining

Finally, we examined subtype distribution across anatomical sites and found that primary pancreatic tumors were more frequently classified as classical, whereas metastatic lesions were enriched for the null subtype (Supplementary Fig. 2c), suggesting molecular evolution during disease progression in PDAC^11^. However, there is no significant association of clinical stage and molecular subtypes. Together, these findings demonstrate that multiplex imaging–based classification using GATA6 and CK5 delineates distinct tumor lineage states in advanced PDAC, with significant associations to patient survival that are not captured by conventional clinical variables.

### Molecular subtypes associate with distinct immune infiltration patterns in advanced PDAC

To determine whether tumor lineage cell states are associated with distinct tumor microenvironment features, we performed multiplex spatial imaging using a panel of markers including lymphoid (CD4, CD8, CD20), myeloid (CD68, CD163), functional (Ki67, FOXP3, GZB, PD1, PDL-1, TIM-3), and structural compartments, including tumor (pan-cytokeratin (PanCK), stromal (SMA), and vascular (CD31) markers (Fig. 2a; Supplementary Fig.3a). Using supervised cell phenotyping, we resolved eighteen distinct cellular populations according to marker expression (Figure.2b). These included major immune subsets such as CD8⁺ T cells—along with functionally defined T cell states (CD8⁺GZB⁺, CD8⁺Ki67⁺, CD8⁺PD-1⁺, CD8⁺TIM-3⁺, CD8⁺PDL-1⁺), CD4⁺T cells, CD4⁺TIM-3⁺, CD4⁺FOXP3⁺. Myeloid populations comprised M1 (CD68⁺CD163⁻) and M2 (CD68⁺CD163⁺) macrophages, as well as PDL1–expressing subsets (CD68⁺PDL-1⁺ and CD163⁺PDL-1⁺). In addition, we identified cancer-associated fibroblasts (CAFs), endothelial cells, B cells, and proliferative or tumor cell subsets (PK⁺Ki67⁺, PK⁺PDL-1⁺). Rule of defining functional cell phenotype was briefly described in Table 2.

**Fig. 2.**
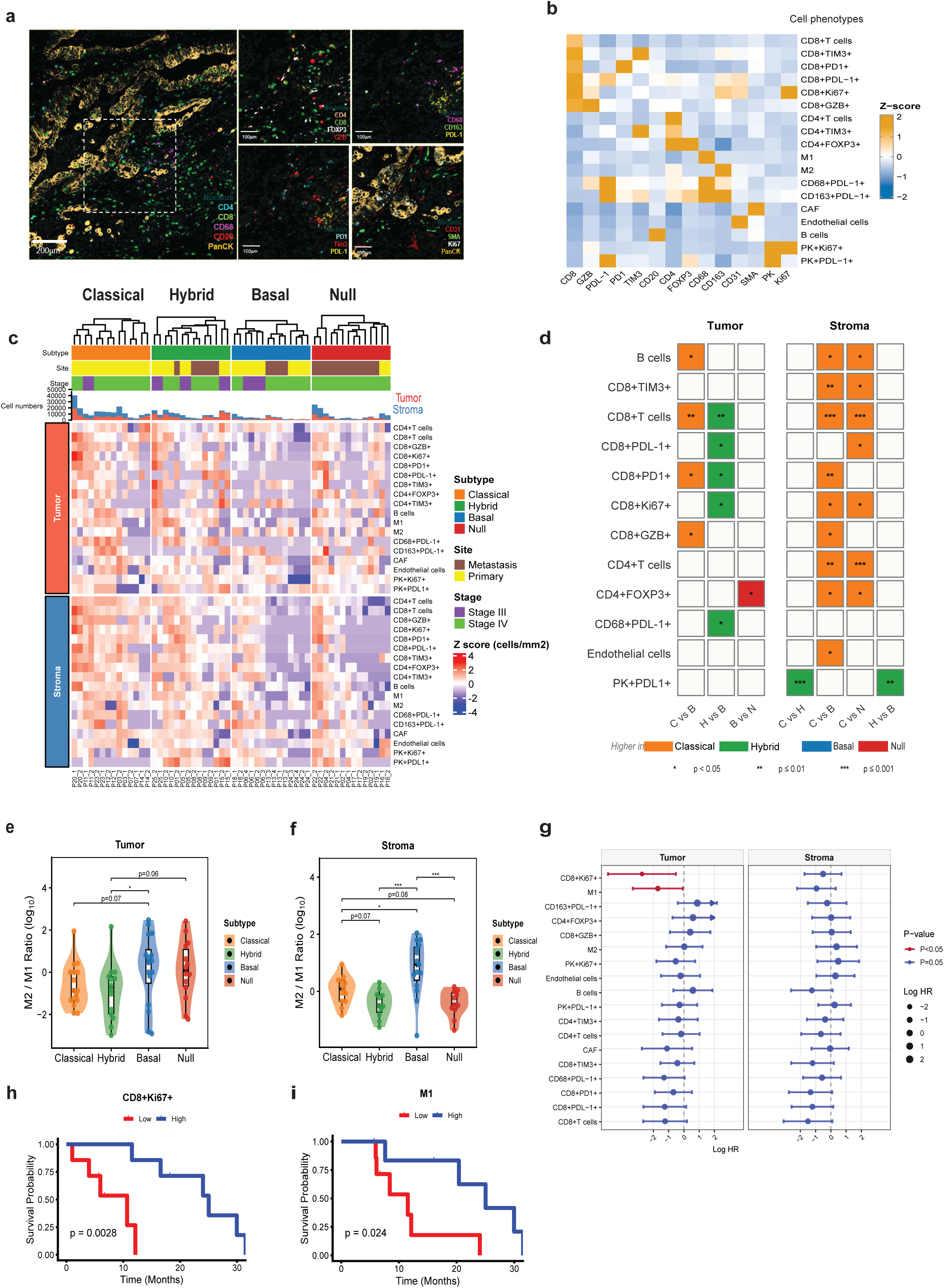
Molecular Subtypes Exhibit Distinct Tumor Immune Microenvironment Architectures. **a** Representative multiplex immunofluorescence images of four PDAC molecular subtypes showing lymphoid markers (CD4, CD8, CD20), myeloid markers (CD68, CD163), functional markers (Ki67, FOXP3, GZB, PDL-1), and structural markers (PK, SMA, CD31). Scale bars, 100-200 μm. **b** Heatmap of marker expression illustrating z-scored marker intensities across individual cellular phenotypes. **c** Heatmap displaying z-score normalized cell densities for immune and stromal cell phenotypes per ROI (column) across molecular subtypes in tumor and stromal compartments. Patients are clustered by subtype. Color scale represents z-score normalized cell density (cells/mm²). **d** Heatmap displaying significant pairwise differences in cell type densities between molecular subtypes (Classical [C], Hybrid [H], Basal [B], and Null [N]) across tumor and stromal compartments. Each cell represents a significant pairwise comparison (Dunn’s post-hoc test following Kruskal-Wallis; *p < 0.05, **p ≤ 0.01, ***p ≤ 0.001). Cell color indicates the subtype with significantly higher density in each comparison: blue = Basal, orange = Classical, green = Hybrid, red = Null. Empty cells indicate non-significant comparisons. **e-f** M2/M1 ratios across molecular subtypes in tumor and stroma compartments. Statistical significance was assessed by Kruskal-Wallis’s test with post-hoc pairwise comparisons. *p < 0.05, **p ≤ 0.01, ***p ≤ 0.001. **g** Forest plot of univariate Cox proportional hazards regression for immune cell phenotypes in the tumor compartment. Log HR and 95% confidence intervals are shown. **h-i** Kaplan–Meier overall survival curves stratifying patients by upper and lower interquartile CD8^+^Ki67^+^ and M1 macrophage cell density. Statistical significance assessed by log-rank test; p values are indicated. ROI= region of interest. All analysis was done in all patients (n=27).

**Table 2.** Cell phenotype classification and marker combinations used for multiplex immunofluorescence phenotyping.

| No. | Cell Phenotype (name) | Lineage | Marker Combination | Functional cell phenotype |
| --- | --- | --- | --- | --- |
| 1 | CD4+T cells | Lymphoid | CD4+CD8- | Helper T cells |
| 2 | CD8+T cells | Lymphoid | CD8+CD4- | Cytotoxic T cells |
| 3 | CD8+GZB+ | Lymphoid | CD8+GZB+ | Activated cytotoxic T cells |
| 4 | CD8+Ki67+ | Lymphoid | CD8+Ki67+ | Proliferating cytotoxic T cells |
| 5 | CD8+PD1+ | Lymphoid | CD8+PD1+ | PD1+ CD8+ T cells |
| 6 | CD8+PDL-1+ | Lymphoid | CD8+PD-L1+ | PDL-1-expressing CD8+ T cells |
| 7 | CD8+TIM-3+ | Lymphoid | CD8+TIM-3+ | TIM-3+ CD8+ T cells |
| 8 | CD4+FOXP3+ | Lymphoid | CD4+FOXP3+ | Regulatory T cells |
| 9 | CD4+TIM-3+ | Lymphoid | CD4+TIM-3+ | TIM-3+ CD4+ T cells |
| 10 | B cells | Lymphoid | CD20+ | B cells |
| 11 | M1 | Myeloid | CD68+CD163- | M1 macrophages |
| 12 | M2 | Myeloid | CD68+CD163+ | M2 macrophages |
| 13 | CD68+PDL-1+ | Myeloid | CD68+PDL-1+ | PDL-1-expressing M1-like macrophages |
| 14 | CD163+PDL-1+ | Myeloid | CD163+PDL-1+ | PDL-1-expressing M2-like macrophages |
| 15 | CAF | Stromal | α-SMA+ | Cancer-associated fibroblasts |
| 16 | Endothelial cells | Stromal | CD31+ | Endothelial cells |
| 17 | PK+Ki67+ | Tumor | PK+Ki67+ | Proliferating tumor cells |
| 18 | PK+PD-L1+ | Tumor | PK+PDL-1+ | PD-L1-expressing tumor cells |

The area-normalized cell density of each cell type was measured in the tumor and stroma regions for each ROI from individual patient. Hierarchical clustering with each subtype group revealed that the immune cell composition within the tumor compartment was broadly comparable across the four subtypes. In contrast, stromal regions of the Basal and Null subtypes exhibited reduced immune cell enrichment relative to the Classical and Hybrid subtypes (Fig. 2c). We further performed post hoc pairwise comparisons to evaluate cell density variation between subtypes (Fig. 2d). In the tumor compartment, the Classical subtype exhibited significantly higher densities of B cells, CD8⁺ T cells, CD8⁺PD1⁺ and CD8⁺GZB⁺ cells compared with the Basal subtype (p < 0.05-0.01). Similarly, the Hybrid subtype demonstrated significantly elevated densities of CD8⁺ T cells, CD8⁺PDL-1⁺, CD8⁺PD1⁺, and CD8⁺Ki67⁺ cells relative to the Basal subtype (p < 0.05-0.01). Additionally, the Classical subtype showed significantly greater stromal CD8⁺ T cell (p ≤ 0.001) and CD4^+^ cell densities (p ≤ 0.001) compared with the Null subtype (Fig. 2d). CD68⁺PDL-1⁺ cells were increased only in the Hybrid subtype. Overall, both Basal and Null subtypes were consistently depleted of cytotoxic, exhausted, and helper T cell populations across both tumor and stromal compartments (Fig. 2d). These findings suggest that the enrichment of lymphoid cells was higher in the Classical and Hybrid subtypes (immune rich phenotypes) whereas the basal and null subtypes exhibited as the immune excluded phenotypes.

We next examined the balance between anti-tumor and immunosuppressive immune responses by calculating the ratios of M2 to M1 macrophages and regulatory T cells to CD8⁺ T cells across tumor subtypes^24, 25^. Notably, the M2:M1 macrophage ratios were significantly elevated in the Basal subtype compared with the Hybrid subtype within the tumor compartment (p < 0.05; Fig. 3e). Consistently in the stroma area, the M2:M1 ratios were also higher in the Basal subtype compared with the Null (p ≤ 0.001), the Hybrid (p ≤ 0.001), and Classical subtypes (p < 0.05) (Fig. 3f). Higher Tregs/CD8 ratios were also found in Basal and Null versus Hybrid (p < 0.05) in tumor area (Supplementary Fig.3b-c). These findings indicate that Basal and null tumors are characterized by a more immunosuppressive microenvironment driven by M2 macrophages and regulatory T cells.

**Fig. 3.**
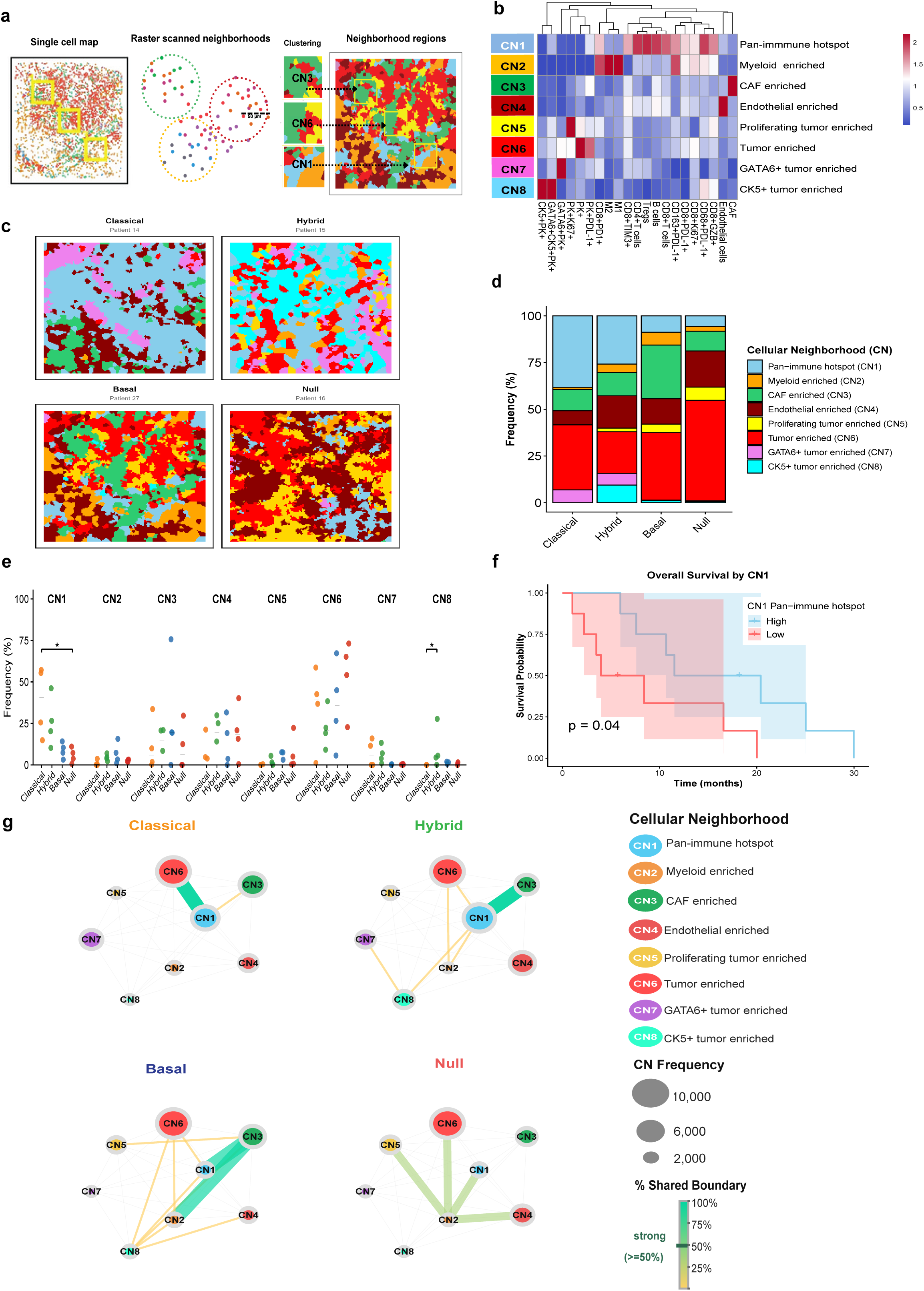
Spatial cellular neighborhood analysis reveals distinct TME organization across PDAC molecular subtypes. **a** Schematic overview of the raster-scanned neighborhood analysis pipeline using Cytomap. **b** Heatmap showing the fold change enrichment of cell types across the eight identified cellular neighborhoods (CN1–CN8). Color intensity represents scaled cell type frequency. Rows represent cellular neighborhoods; columns represent cell types; Pan-immune hotspot (CN1), Myeloid enriched (CN2), CAF enriched (CN3), Endothelial enriched (CN4), Proliferating tumor enriched (CN5), Tumor enriched (CN6), GATA6+ tumor enriched (CN7), and CK5+ tumor enriched (CN8). **c** Spatial map of representative fields of view (FOVs) from classical, hybrid, basal and null subtype tumors, with cells colored by assigned cellular neighborhoods in (b). **d** Stacked bar plots showing the mean proportion of CN composition (CN1–CN8) in each subtype. Each color represents a distinct CN. **e** Box plots comparing the prevalence (%) of each cellular neighborhood (CN1–CN8) across molecular subtypes (classical, hybrid, basal, null). Differences across subtypes were assessed using Kruskal-Wallis’s test with post-hoc pairwise comparisons. *p < 0.05, **p ≤ 0.01, ***p ≤ 0.001. **f** Kaplan–Meier overall survival curves stratified Pan-immune hotspot neighborhoods (CN1) frequencies, dichotomized at the median value. Survival analysis was done using Kaplan-Meier method with log-rank test. **g** CN–CN spatial interaction networks for Classical, Hybrid, Basal, and Null subtypes. Nodes represent cellular neighborhoods, sized proportionally to CN abundance. Edge thickness reflects the frequency of shared boundaries between CN pairs.

Given the distinct landscape observed in different tumor subtypes, we next evaluated whether the cellular abundance was associated with patient outcomes using univariate Cox proportional hazards regression across all 18 cell types in both tumor and stromal compartments (Fig. 2g). Among these, CD8⁺Ki67⁺, proliferating CD8 T cells (Log HR = −2.81, 95% CI: −4.61 to −0.53, p = 0.015) and M1macrophages (Log HR = −1.71, 95% CI: −3.22 to −0.07, p = 0.041) were significantly associated with improved survival. Consistent with the observed enrichment of M2 macrophages in Basal tumors, these findings further highlight the prognostic importance of macrophage polarization, where a shift toward an M1-dominant state is linked to favorable outcomes. Accordingly, higher densities of CD8⁺Ki67⁺ T cells and M1 macrophages were associated with prolonged median survival (25.0 vs 10.7 months, p = 0.0028; Fig. 2h and 24.7 vs 11.4 months, p = 0.024, respectively; Fig. 2i), supporting a role for active immune effector function in driving favorable outcomes, in line with prior studies ^26, 27^. These results collectively link subtype-specific immune architecture to clinical outcomes.

### Spatial neighborhood analysis reveals subtype-specific cellular organization in the PDAC tumor microenvironment

Building on the observed subtype-specific differences in immune composition and their prognostic relevance, we next investigated the spatial organization of the tumor microenvironment which has been shown to reflect underlying immune responses^28^. Raster-scanned neighborhood analysis was assigned to quantify local cellular composition using CytoMAP^29^ (n=16) (Fig. 3a). This approach identified eight distinct cellular neighborhoods (CNs), each defined by dominant cell-type enrichment: Pan-immune hotspot (CN1), Myeloid enriched (CN2), CAF enriched (CN3), Endothelial enriched (CN4), Proliferating tumor enriched (CN5), Tumor enriched (CN6), GATA6⁺ tumor enriched (CN7), and CK5⁺ tumor enriched (CN8) (Fig. 3b). The four molecular subtypes exhibited distinct cellular neighborhood organization patterns with Classical and Hybrid subtypes were occupied with higher CN1 compartment whereas Null subtype showed higher tumor enriched (CN6) (Fig. 3c-d). We then measured the prevalence frequency of each CN and compared among subtypes; the Classical subtype is highly enriched with CN1 when compared with Null subtype (p < 0.05) whereas CN8 was enriched for the Hybrid compared to the Classical subtype (p < 0.05) (Fig.3e). We divided the patients on CN abundance to evaluate the clinical relevance of spatial organization. Univariable Cox proportional hazards and survival analyses showed that higher CN1 abundance was associated with improved overall survival (log-rank p = 0.04; Fig. 3f), with a hazard ratio (HR) of 0.29 (95% CI: 0.08–1.01, p = 0.052; Supplementary Fig. 4b). In contrast, higher CN6 abundance was associated with poorer clinical outcome (log-rank p = 0.078), with an HR of 2.66 (95% CI: 0.86–8.24, p = 0.090; Supplementary Fig. 4a-b). These results also confirm that cellular organization like pan-immune hotspot (CN1) in Classical and Hybrid subtype may enhance the anti-tumor response rather than individual immune cells; leading to have lower tumor enriched (CN6) in these subtypes.

To capture broader tissue architecture, we mapped how cellular neighborhoods relate to one another by constructing CN–CN interaction networks where each node’s size reflects how frequently a neighborhood appears, and each edge’s thickness encodes how often two neighborhoods share a physical boundary across the patient cohort (Fig. 3g). Classical tumors exhibited prominent CN1–CN6 interactions, suggesting increased spatial integration between immune and tumor compartments. Hybrid tumors displayed CN1–CN3 interactions, indicating partial immune engagement within a stromal context. In contrast, basal tumors were characterized by higher CAF enriched (CN3) which act as a bridging node between Pan-immune hotspot (CN1) and Myeloid enriched (CN2) neighborhoods. Null tumors also showed a myeloid-dominant architecture, in which CN2 served as the central and connecting to the Tumor enriched (CN6), Endothelial enriched (CN4), and CAF enriched (CN3) neighborhoods (Fig. 3g). These findings support that distinct spatial cellular organization in the Classical and Hybrid tumors exhibit as immune-accessible architectures, whereas basal and null tumors are characterized by stromal and myeloid dominated networks that may constrain effective tumor–immune interactions.

### Spatial positioning of cytotoxic CD8⁺ T cells relative to helper CD4⁺ T cells and suppressive macrophages predict patient survival

After we evaluated distinct spatial cellular neighborhoods among subtypes, we hypothesized that spatial relationships between immune populations rather than their abundance alone might display more spatial immune architecture insights. Here, we established a spatial interaction scoring framework integrating radius and k-nearest neighbor-based approaches to quantify pairwise cell–cell interactions (Fig. 4a; see details in Methods). Overall, the immune cells preferentially interacted with one another rather than with tumor epithelial cells, indicating spatial segregation between immune and tumor compartments in PDAC tumor microenvironment (Supplementary Fig. 5a-b).

**Fig. 4.**
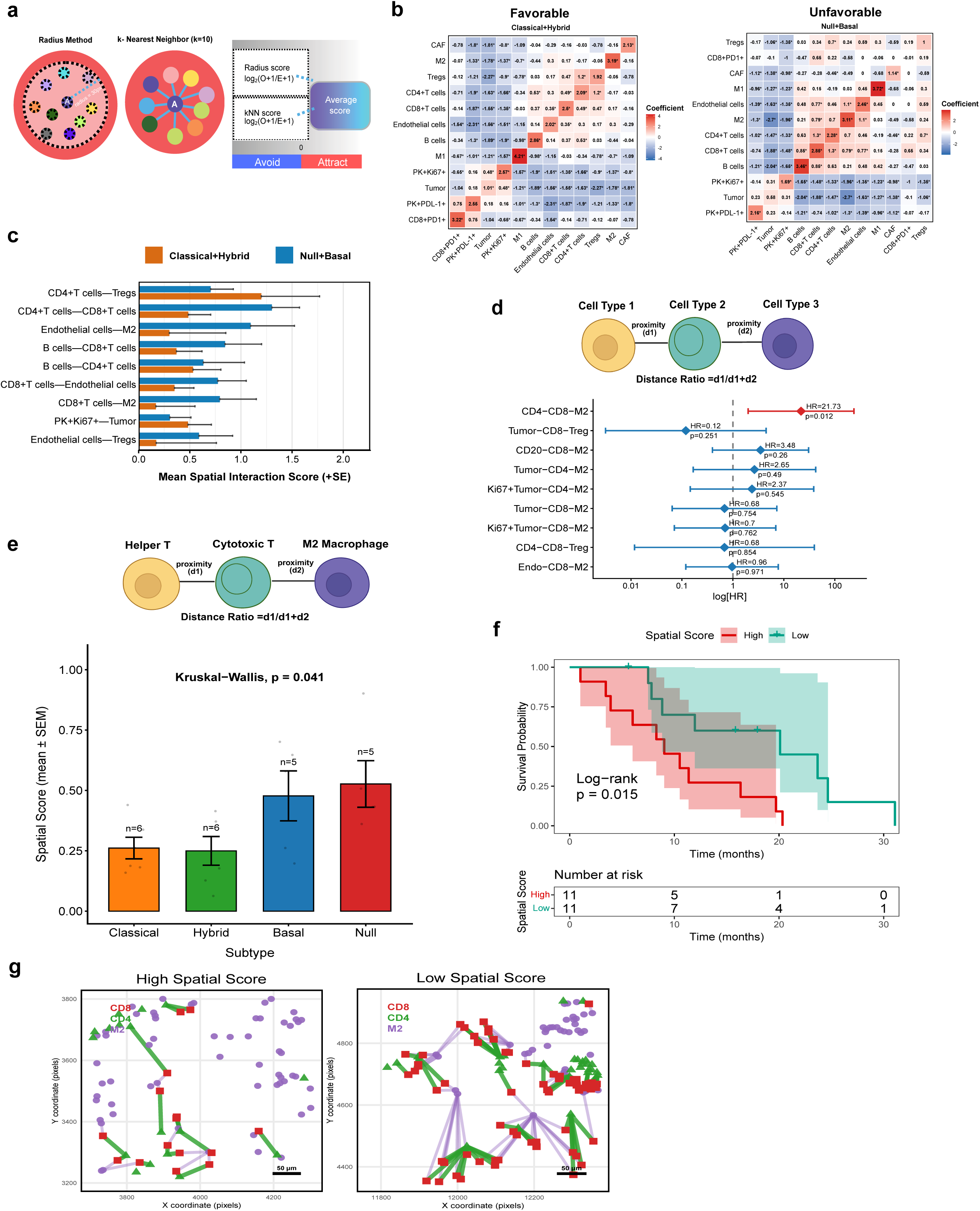
Spatial cell-cell interaction analysis revealed a unique spatial positioning associated with clinical outcome. **a** Workflow of spatial cell-cell interaction analysis; implementing integrated radius and KNN methods. **b** Spatial interaction heatmaps stratified by Classical+Hybrid (n=8) and Null+Basal (n=8) subgroups in selected cell types (tumor, immune, stroma). Positive scores indicate spatial co-localization; negative scores indicate spatial avoidance. Asterisks denote statistically significant pairs. **c** Bar plot of mean interaction scores for significant cell–cell pairs in Classical+Hybrid (orange) (n=8) and Null+Basal (blue) (n=8) subtypes. Bar plots display Mean interaction scores (+SEM) for statistically significant positive pairs within each subgroup. Classical+Hybrid (orange) and Null+Basal (blue) are shown side by side for visual comparison. **d** Schematic of the spatial score framework; showing position of cell types in spatial score formulation. Forest plot below shows univariate Cox regression results (log-scale HR and log-rank p) for nine candidate scores evaluated. **e** Bar plots of spatial scores across classical, hybrid, basal, and null molecular subtypes. Error bars represent standard error of the mean. **f** Kaplan–Meier overall survival curves stratified by spatial score (high, red; low, green; dichotomized at median). **g** Representative spatial maps of FOVs with high (left) and low (right) spatial scores. Each connecting line represents the distance between CD8 to nearest CD4 or M2 macrophage. b: Color intensity reflects spatial interaction scores (red, high; blue, low/avoidance) and statistical significance was described as *p < 0.05, **p < 0.01, ***p < 0.001. FOV= field of view.

Tumors were then stratified into favorable (Classical and Hybrid) and unfavorable (Basal and Null) subtypes based on established PDAC subtype classifications and observed survival patterns where favorable subtypes (Classical and Hybrid) demonstrated significantly longer overall survival compared with unfavorable subtypes (Basal and Null) (log-rank p = 0.0025), with median overall survival of 20.11 months versus 8.28 months, respectively (Supplementary Fig. 1b). Favorable subtypes (classical and hybrid) showed enriched interactions between CD8⁺ T cells with endothelial cells and CD4⁺ T cells (Fig. 4b-c), consistent with coordinated immune activation and trafficking. Notably, unfavorable subtypes (basal and null) exhibited increased interactions between M2 macrophages and both CD8⁺ T cells and endothelial cells (Fig.4b), suggesting immunosuppressive microenvironment that may limit effective cytotoxic T cell recruitment or function^30^. We hypothesized that pairwise interactions do not capture how CD8⁺ T cells are positioned within competing immune niches, we first selected the significant positive pairwise interactions within each clinical-outcome stratified group (Fig.4c). We next constructed triadic spatial positioning scores based on distance ratios among anchor, target, and suppressor populations. Using CD8⁺ T cells as the anchor, we defined nine candidate scores incorporating supportive targets (CD4⁺ T cells, tumor cells, B cells, endothelial cells) and suppressive populations (CD163⁺ M2 macrophages and regulatory T cells) (Fig. 4d). Each score quantified the relative positioning of CD8⁺ T cells as the ratio of distance to the target population over the combined distance to both target and suppressor populations. Among the nine-candidate metrics, only the CD4–CD8–M2 triadic score was significantly associated with overall survival (HR =21.73, p = 0.012), indicating that CD8⁺ T cells positioned closer to suppressive M2 macrophages than to supportive CD4⁺ T cells confer a substantially elevated risk of death (Fig. 4d). Notably, the Basal and Null tumors exhibited significantly higher scores compared with classical and hybrid tumors (Fig. 4e), indicating a shift of CD8⁺ T cells toward suppressive niches in unfavorable states. Survival analysis confirmed that patients with high CD4–CD8–M2 scores had significantly worse overall survival than those with low scores (Fig. 4f), with median survival of 9.03 versus 20.11 months. Representative spatial maps further illustrate that high-score tumors are characterized by CD8⁺ T cells positioned more distant from CD4⁺ T cells and closer to M2 macrophages (Fig.4g). Notably, spatial proximity between CD8⁺ T cells and tumor cells alone was not associated with survival in our cohort (Fig. 4d), supporting that the prognostic impact of CD8⁺ T cells in advanced PDAC were determined by their spatial positioning within competing immune niches, specifically, their relative proximity to supportive CD4⁺ helper T cells versus suppressive M2 macrophages.

### Hybrid PDAC tumors exhibit spatially segregated epithelial states with distinct immune microenvironments

Intratumoral heterogeneity was evident across PDAC subtypes, with hybrid tumors uniquely enriched for epithelial cells co-expressing GATA6 and CK5^10, 12, 23^. To determine whether this tumor cell lineage diversity is associated with spatially distinct tumor microenvironments, we assessed hybrid tumors (n=4) with sufficient tissue architecture to assess intratumoral heterogeneity of tumor cells. Interestingly, spatial mapping revealed that classical (GATA6⁺CK5⁻PK⁺) and basal (GATA6⁻CK5⁺PK⁺) tumor cells occupied discrete regions of higher classical and higher basal regions within an individual tumor (Fig.5a), demonstrating pronounced spatial segregation of epithelial states at single-cell resolution (Fig.5b). These observations are consistent with prior transcriptomic studies describing intermediate epithelial states co-expressing classical and basal lineage programs^13^.

To further investigate whether spatially segregated epithelial states are associated with distinct local immune microenvironments, tumor regions were stratified into classical-enriched (Hybrid_Classical) and basal-enriched (Hybrid_Basal) cores based on regional tumor cell type composition (Fig. 5c). Mean threshold of proportion of each cell type (Basal and Classical cells) was used for classifying the Hybrid_Basal and Hybrid_classical cores in Hybrid tumors. To determine whether these spatially segregated epithelial states were associated with distinct local immune microenvironments, we quantified area-normalized immune cell densities within Hybrid_Classical and Hybrid_Basal regions. Although CD8⁺ T cells tended to be more abundant in Hybrid_Classical regions and Tregs showed a trend toward enrichment in Hybrid_Basal regions, these differences did not reach statistical significance (Fig. 5d). In contrast, M2 macrophages were significantly enriched in Hybrid_Basal cores (p = 0.013; Fig. 5d), suggesting the presence of a more immunosuppressive niche.

**Fig. 5.**
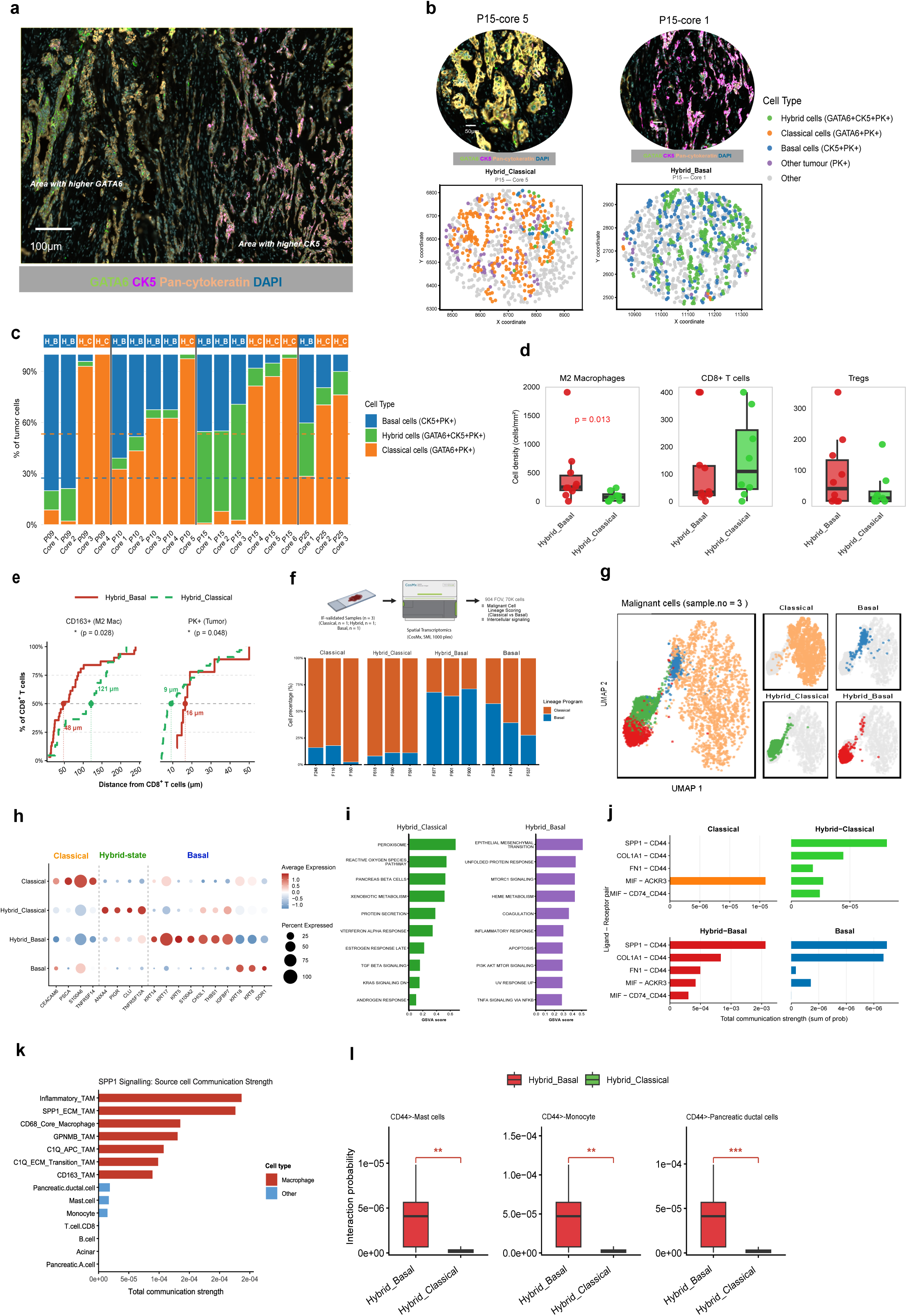
Spatial immune heterogeneity and lineage–immune coupling in hybrid pancreatic tumors. **a** Representative multiplex immunofluorescence images showing spatial distribution of GATA6 (green), CK5 (magenta), and Pan-cytokeratin (orange) with DAPI counterstain (blue) within a hybrid PDAC tumor. Scale bar = 100 µm. **b** Representative multiplex immunofluorescence image (upper panel) with single spatial map (lower panel) showing core region (P15-core5) having higher GATA6 (green) and region (P15_core1) having higher CK5 (magenta) with DAPI counterstain (blue) within a hybrid PDAC tumor (P15). Scale bar = 50 µm. Lower panel illustrating the subtype specific cell types. **c** Stacked bar plots showing the proportion of epithelial lineage states classical (GATA6⁺PK^+^), basal (CK5⁺PK^+^), and hybrid (GATA6⁺CK5⁺PK^+^) across individual tumor cores. A threshold based on mean proportion across the tumor was used to stratify tumor cores into lineage-enriched groups (Hybrid–Classical and Hybrid–Basal). Blue and orange dot lines represent for each threshold for classical and basal enriched regions respectively. **d** Comparison of area-normalized densities of M2 macrophages, CD8⁺ T cells and Tregs across Hybrid–Classical (n = 4) and Hybrid–Basal (n = 4). Statistical comparisons by Wilcoxon rank-sum test. **e** Spatial distance analysis showing cumulative distance of CD8⁺ T cells relative to CD163⁺ macrophages and tumor epithelial cells across Hybrid–Classical and Hybrid–Basal tumors. **f** Workflow of spatial transcriptomics 1K panel: Upper panel. Stacked bar plots showing the relative proportion of Classical and Basal transcriptional scores per FOV, derived from gene signatures, used to assign FOV-level lineage identity across three PDAC samples profiled: Lower panel. **g** UMAP visualization of malignant ductal cells from CosMx spatial transcriptomics (n = 3 patients; Classical, Hybrid, Basal), annotated by transcriptional lineage groups from FOVs based classification. The inset shows proportional distribution of clusters across Basal, Classical, and Hybrid_Classical and Hybrid_Basal. **h** Expression of lineage-associated marker genes across FOVs classified lineage groups, validating cell states. **i** GSVA pathway enrichment across malignant lineage groups, highlighting distinct biological programs. **j** Top ligand-receptor signaling interactions across PDAC subtypes ranked by total communication strength (sum of interaction probabilities). **k** Bar plots showing total outgoing SPP1 ligand communication strength per source cell type, summed across all ligand–receptor interactions from different cell types (Macrophage, red) and non-macrophage populations (Other, blue) distinguished by color. Cell types ranked by descending communication strength. **l** Box plots showing inferred SPP1–CD44 ligand–receptor interaction strength across target cell populations among tumor lineage groups, highlighting increased signaling in Hybrid–Basal regions.

Because immune cell abundance alone may not fully capture functional immune organization, we next examined the spatial positioning of CD8⁺ T cells relative to tumor cells and M2 macrophages. CD8⁺ T cells in Hybrid_Classical regions were positioned closer to tumor epithelial cells (9 µm) and farther from M2 macrophages (121 µm), whereas CD8⁺ T cells in Hybrid_Basal regions were located farther from tumor cells (48 µm) and in closer proximity to M2 macrophages (16 µm) (p < 0.05; Fig. 5e). These findings indicate that Hybrid_Classical regions exhibit a more immune-engaged spatial configuration, whereas Hybrid_Basal regions are characterized by a macrophage-associated suppressive niche. Together, these results suggest that local epithelial lineage states are linked not only to immune cell composition but also to the spatial organization of the immune microenvironment within hybrid tumors.

### Hybrid tumor states exhibit distinct transcriptional programs and macrophage-derived signaling networks

Having established that hybrid PDAC tumors harbor spatially segregated epithelial states associated with distinct immune microenvironments at the protein level, we next sought to resolve the transcriptional basis and intercellular signaling mechanisms underlying these observations. We performed CosMx 1000-plex spatial transcriptomics on three PDAC samples. Each subtype category had been validated by the t-CyCIF experiment (Fig. 5f, upper panel). This approach enabled simultaneous single-cell transcriptome profiling while preserving spatial coordinates at subcellular resolution. To systematically capture epithelial lineage heterogeneity at the level of tumor microregions, mirroring the protein level region-wise stratification, we applied a field-of-view (FOV)–level classification framework on 904 FOVs using the transcriptional classical and basal programs of constituent malignant cells. FOVs from molecularly classified Classical or Basal tumors were assigned according to mIF, while FOVs from Hybrid tumors were subdivided by the Delta score directionality (Hybrid_Classical: Delta ≥ 0.1; Hybrid_Basal: Delta < 0.1) (see more details in Methods). FOVs with missing subtype annotations were excluded. With this approach, we can stratify tumor regions into four epithelial lineage groups, namely Classical, Basal, Hybrid_Classical and Hybrid_Basal from three samples of CosMx data (Fig.5f: lower panel). To assess whether the spatially defined FOV lineage groups represented distinct transcriptional states, we performed supervised UMAP analysis of malignant cells across the three CosMx samples. Four well-separated clusters emerged, corresponding to the Classical, Hybrid_Classical, Hybrid_Basal, and Basal FOV-level groups, indicating that the spatial classification captures biologically distinct malignant cell states (Fig. 5g). Within these lineage groups, Hybrid_Classical cells were characterized by enrichment of Hybrid-state markers, including ANXA4, PIGR, CLU, and TNFRSF12A, whereas Hybrid_Basal cells exhibited increased expression of Basal-associated genes such as KRT14, KRT17 and S100A2 together with CHI3L1, THBS1, and IGFBP7 (Fig. 5h; Supplementary Fig. 6a). Classical markers (CEACAM6, PSCA, S100A6, and TNFRSF14) remained predominantly restricted to the Classical group, while Basal markers were progressively acquired across Hybrid_Basal and Basal states. These expression patterns indicate that Hybrid_Classical and Hybrid_Basal represent transcriptionally distinct states within the Classical–Basal lineage spectrum. Hybrid_Classical was characterized by enrichment of Hybrid-state markers, whereas Hybrid_Basal showed acquisition of Basal-associated genes together with Hybrid-state features. Together, these findings suggest that the Hybrid subtype comprises multiple molecularly distinct epithelial states rather than a single homogeneous Hybrid population.

Consistent with the distinct transcriptional identities of the two Hybrid states, GSVA revealed marked functional divergence between Hybrid_Classical and Hybrid_Basal regions (Fig. 5i). Hybrid_Classical was enriched for immune-responsive and metabolic programs, including interferon-α response, reactive oxygen species signaling, peroxisome activity, and protein secretion. In contrast, Hybrid_Basal showed enrichment of epithelial–mesenchymal transition, mTORC1 signaling, inflammatory TNFα signaling via NF-κB, unfolded protein response, and PI3K–AKT–mTOR signaling pathways. These findings indicate that the two Hybrid states are associated with distinct biological programs, with Hybrid_Basal exhibiting a more inflammatory and dedifferentiated transcriptional phenotype relative to Hybrid_Classical.

To investigate the intercellular signaling mechanisms associated with lineage-specific immune niches, we applied CellChat to the spatial transcriptomic dataset. Co-registration of 19,118 macrophages to the epithelial lineage classification framework followed by unsupervised clustering identified seven transcriptionally distinct macrophage populations, including SPP1/ECM TAMs, C1Q/APC TAMs, Inflammatory TAMs, C1Q/ECM Transition TAMs, CD68 Core Macrophages, CD163 TAMs, and GPNMB TAMs (Supplementary Fig. 6b). Marker gene expression patterns supported the annotation of these macrophage populations, including SPP1 and APOE expression in SPP1/ECM TAMs, CXCL8 and IL1B expression in Inflammatory TAMs, C1Q-family gene expression in C1Q/APC and C1Q/ECM Transition TAMs, CD68 in Core Macrophages, CD163 expression in CD163 TAMs, and GPNMB expression in GPNMB TAMs (Supplementary Fig. 6c). Although SPP1-associated macrophage populations, including SPP1/ECM TAMs and Inflammatory TAMs, were observed across all epithelial lineage groups (Supplementary Fig. 6d), we hypothesized that their signaling activity might differ according to epithelial state. CellChat analysis identified SPP1–CD44 as the dominant ligand-receptor interaction associated with Basal-lineage tumor states. SPP1–CD44 signaling was absent in Classical regions, emerged in Hybrid_Classical regions, and reached its highest communication strength in Hybrid_Basal regions before persisting in Basal tumors (Fig. 5j). In contrast, other ligand-receptor interactions, including COL1A1–CD44, FN1–CD44, MIF–ACKR3, and MIF–CD74/CD44, exhibited substantially lower communication strength.

To identify the cellular source of SPP1 signaling, we ranked outgoing communication strength across all cell populations. Inflammatory_TAMs and SPP1_ECM_TAMs emerged as the dominant SPP1-secreting populations, far exceeding contributions from other immune and stromal cell types (Fig. 5k). Consistent with this finding, analysis of the broader macrophage communication network identified SPP1–CD44 as the highest-probability macrophage-derived ligand–receptor interaction, surpassing FN1–CD44, MIF–ACKR3, COL1A1–CD44, MIF–CD74/CD44, PECAM1–PECAM1, and SPP1– ITGAV_ITGB5 interactions (Supplementary Fig. 6e). Finally, direct comparison of macrophage-derived SPP1–CD44 signaling between Hybrid_Basal and Hybrid_Classical regions demonstrated significantly greater communication probability in Hybrid_Basal across multiple receiver populations, including mast cells, monocytes, and pancreatic ductal cells (Fig. 5l; Wilcoxon test). Together, these findings identify macrophage-derived SPP1–CD44 signaling as a lineage-state-dependent communication program preferentially activated in Hybrid_Basal tumor microenvironment.

### Integration of spatial immune score and GATA6 expression improves prognostic discrimination in advanced PDAC

To evaluate the prognostic relevance of spatial immune organization in the context of established clinicopathological variables, we performed Cox proportional hazards analysis (Fig. 6a). In univariate analysis, both the spatial score (HR = 1.89, 95% CI 1.15–3.11, p = 0.012) and unfavorable molecular subtype (HR = 4.77, 95% CI 1.55–14.68, p = 0.006) were significantly associated with worse overall survival. In contrast, metastatic stage, CA19-9 level, age, gender, M1 macrophage density, and CD8⁺Ki67⁺ cell density, were not significantly associated with outcome. Although neither the spatial score nor molecular subtype remained statistically significant after adjustment, both retained elevated hazard ratios, suggesting that spatial immune organization and molecular subtype capture complementary prognostic information that may not be fully resolved given the limited sample size and number of events.

**Fig. 6.**
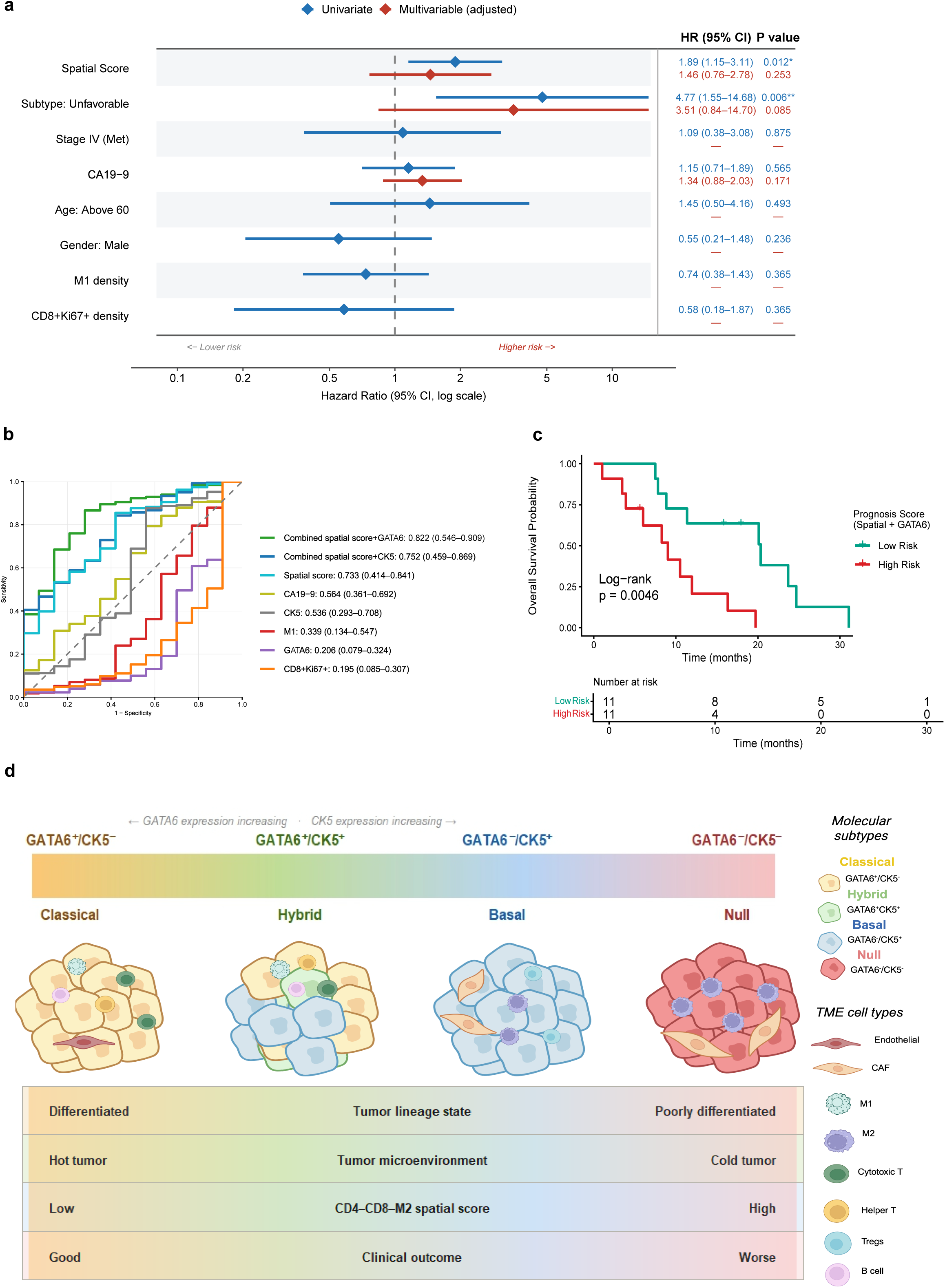
Survival analysis and prognostic integration. **a** Forest plot of univariate and multivariable Cox Proportional Harzard model analyses. Harzard ratios (HR) with 95% confidence intervals are plotted for spatial score, molecular subtypes (Unfavorable), clinical covariates (stage, CA19-9, age, gender), M1 density and CD8+Ki67+ density. HR (95% CI) and P-values are shown on the right. **b** Time-dependent ROC curves with integrated AUC (iAUC) computed using inverse probability of censoring weighting (IPCW) across the follow-up period. Time-dependent ROC curves comparing prognostic discriminatory accuracy of individual markers and combined multivariable models. Data are presented as mean iAUCs values with 95% confidence intervals. **c** Kaplan–Meier survival analysis of a combined prognostic score integrating spatial score and GATA6 expression, dichotomized at the median. Log-rank p-values are shown; p<0.05 was considered statistically significant**. d** Schematic summarizing the integrated molecular and spatial immune landscape across Classical, Hybrid, Basal and Null PDAC subtypes. Gradient bars indicate directional changes in GATA6 expression, CK5 expression, immune composition, spatial score, and clinical outcome across subtypes. This figure was created in https://BioRender.com

To further assess predictive performance, we performed time-dependent receiver operating characteristic (ROC) analysis^31^ (Fig. 6b). Among individual variables, the spatial score demonstrated the strongest prognostic discrimination (AUC = 0.733), outperforming CA19-9 (AUC = 0.564), CK5 expression (AUC = 0.536), M1 macrophage density (AUC = 0.339), GATA6 expression (AUC = 0.206), and CD8⁺Ki67⁺ T-cell density (AUC = 0.195). Notably, integration of the spatial score with GATA6 expression yielded the highest predictive accuracy (AUC = 0.822, 95% CI 0.546–0.909), indicating that spatial immune organization and tumor lineage information provide complementary prognostic value beyond either variable alone. Based on its superior performance, the combined spatial–GATA6 model was used to derive a composite prognostic score. Stratification of patients using the median score identified high- and low-risk groups with significantly different overall survival outcomes (log-rank p = 0.0046; Fig. 6c), supporting the clinical utility of integrating spatial immune context with molecular subtype information.

To summarize the relationship between tumor lineage state, spatial immune organization, and clinical outcome, we developed an integrative model based on our spatial and molecular analyses (Fig. 6d). Classical tumors were characterized by high GATA6 expression, enrichment of CD8⁺ and CD4⁺ T cells, and B cells, together with lower spatial scores, consistent with an immune-enriched microenvironment and favorable clinical outcome. In contrast, Basal and Null tumors exhibited reduced GATA6 expression, increased M2 macrophage predominance, and higher spatial suppression scores, reflecting greater immune exclusion and poorer prognosis. Hybrid tumors generally occupied an intermediate position between these extremes, although spatial transcriptomic analyses revealed substantial heterogeneity within the Hybrid category. Collectively, these findings demonstrate that tumor lineage state and spatial immune organization provide complementary prognostic information in advanced PDAC, and that their integration improves risk stratification beyond conventional molecular classification alone.

## Discussion

Pancreatic ductal adenocarcinoma (PDAC) remains one of the most lethal malignancies, with most patients diagnosed at advanced, unresectable stages. Despite the clinical importance of advanced disease, most molecular and spatial profiling studies have focused on surgically resected tumors, where recent single-cell and spatial analyses have revealed complex immune ecosystems and tumor–stromal interactions^4, 15^. Consequently, the spatial biology of advanced-stage PDAC remains poorly characterized. In this study, we applied highly multiplexed tissue cyclic immunofluorescence (t-CyCIF) to stage III–IV PDAC to investigate how tumor lineage programs interact with the spatial immune microenvironment. In our work, we aimed to connect the tumor differentiation state and spatial immune organization with clinical outcomes in advanced PDAC.

Using multiplexed protein-level profiling, we classified tumors into four lineage states: classical, basal, hybrid, and null, based on GATA6 and CK5 expression. These categories recapitulate molecular frameworks previously established through transcriptomic profiling of PDAC^2, 7, 8^. In our advanced-stage cohort, classical and hybrid tumors were associated with improved overall survival compared with basal and null tumors, consistent with prior reports linking basal-like transcriptional programs with aggressive clinical behavior^23^. Notably, the null subtype emerged as the strongest independent predictor of poor survival in univariable analysis, highlighting a particularly aggressive tumor state that remains incompletely characterized. The spatial immune cell states, together with the coexistence of classical and basal epithelial programs in hybrid tumors, revealed distinct intra-tumoral heterogeneity. These observations align with growing evidence that PDAC cells exhibit substantial transcriptional plasticity and intermediate states along the classical–basal spectrum^11^.

Consistent with some prior studies linking tumor lineage programs to immune contexture, our study identified classical and hybrid tumors that displayed immune-inflamed microenvironments enriched for CD8⁺ cytotoxic T cells, CD4⁺ helper T cells, and B cells. Conversely, basal and null tumors were characterized by immune-excluded phenotypes with reduced lymphocyte infiltration and increased abundance of M2 macrophages. Tumor-associated macrophages are known to suppress cytotoxic T cell activity through multiple mechanisms, including PD-L1 signaling, arginase-mediated metabolic suppression, and secretion of immunomodulatory cytokines such as TGF-β and IL-10^32^. Increasing evidence also indicates that inflammatory signals within the tumor microenvironment can actively drive basal-like transcriptional programs in PDAC cells, highlighting dynamic interactions between tumor identity and immune composition ^11, 19, 20, 33^. Our neighborhood analysis highlighted that Classical tumor exhibited prominent CN1–CN6 interactions, consistent with close spatial coupling between immune and tumor compartments^34^. Consistent with the co-existence of immune-enriched and fibroblast-dominant multicellular communities previously identified in PDAC^35^, Hybrid tumors exhibited interactions between Pan-immune hotspot (CN1) and CAF enriched (CN3) neighborhoods, suggesting a spatially permissive niche for partial immune engagement within a stromal context. Basal subtypes were characterized by CAF–myeloid networks (CN3), consistent with reports that CAFs actively recruit and polarize immunosuppressive myeloid populations in PDAC, establishing physical and biochemical barriers to T-cell infiltration^36^. Interestingly, Null tumors exhibited a distinct immune architecture characterized by myeloid-dominant networks with minimal lymphocyte engagement. These findings suggest that Null tumors represent a distinct immunosuppressed state that differs from the stromal-associated exclusion observed in Basal tumors^33^.

Spatial proteomic and transcriptomic analysis revealed that Hybrid PDAC tumors are not immunologically homogeneous but contain spatially segregated epithelial states associated with distinct immune niches, wherein Classical-enriched regions exhibited immune-permissive architectures and Basal-enriched regions showed increased M2 macrophage density. These findings indicate that epithelial lineage state and local immune architecture are closely linked, consistent with dynamic interactions between tumor cell programs and the surrounding microenvironment. This is consistent with previous studies showing that inflammatory signals within the tumor microenvironment can influence epithelial plasticity and promote basal-like transcriptional programs in PDAC^13^. A notable finding from our spatial transcriptomic analysis was that Hybrid tumors were not simply homogeneous intermediate states but instead contained spatially segregated Hybrid_Classical and Hybrid_Basal regions with distinct transcriptional programs, immune architectures, and cell–cell communication networks. These observations suggest that epithelial plasticity in PDAC occurs at the level of local tumor microregions and may contribute to intratumoral variability in immune engagement and therapeutic vulnerability. Additionally, the coexistence of immune-permissive and immune-suppressive niches within Hybrid tumors may partially explain the intermediate clinical behavior observed for this subtype and highlights the importance of considering intratumoral spatial heterogeneity beyond bulk lineage classification.

Beyond canonical immunosuppressive mechanisms, SPP1⁺ tumor-associated macrophages (TAMs) have been implicated in regulating epithelial plasticity through the SPP1–CD44 signaling axis^37,38^. Consistent with this concept, ligand–receptor analysis identified SPP1–CD44 as the dominant communication pathway in dedifferentiated tumor states, with the strongest signaling detected in Hybrid_Basal regions. Inflammatory TAMs and SPP1/ECM TAMs emerged as the primary cellular sources of SPP1 and were preferentially associated with macrophage-rich, dedifferentiated tumor niches. The selective enrichment of SPP1–CD44 signaling in Hybrid_Basal regions, together with increased M2 macrophage abundance and a more dedifferentiated epithelial phenotype, suggests a potential association between macrophage-derived signaling, epithelial plasticity, and local immune remodeling. These findings are consistent with previous reports implicating SPP1 signaling in cancer stemness and tumor cell plasticity in PDAC^38, 39, 40^ and raise the possibility that epithelial lineage state and macrophage-mediated signaling may reinforce one another during tumor progression.

Beyond immune composition alone, our study highlights the importance of spatial immune organization in determining clinical outcomes. Increasing evidence across multiple cancer types suggests that spatial relationships among immune populations provide more prognostic information than immune cell abundance alone ^28, 41^. Consistent with this concept, conventional abundance-based immune metrics, including CD8⁺Ki67⁺ cell densities and M1 macrophage cell densities, did not reach the highest prognosis value in our cohort, whereas the spatial score emerged as one of the strongest prognostic variables. These findings suggest that the functional state of the tumor immune microenvironment is determined not only by the presence of immune cells but also by their spatial positioning relative to one another.

By systematically evaluating candidate spatial metrics, we identified a spatial score that captures the relative proximity of CD8⁺ cytotoxic T cells to CD4⁺ helper T cells versus CD163⁺ immunosuppressive macrophages. Higher scores were associated with poorer clinical outcome, indicating that unfavorable tumor states are characterized by CD8⁺ T cells positioned within macrophage-dominated suppressive niches rather than immune-supportive microenvironments. These observations extend earlier studies linking CD8⁺ T cells infiltration with prognosis in PDAC^42^ by demonstrating that spatial context provides additional biological and prognostic information beyond immune cell abundance alone. Importantly, integration of the spatial score with GATA6 expression yielded the highest prognostic discrimination in our cohort, substantially outperforming either variable alone. This finding suggests that GATA6 captures tumor-intrinsic differentiation state, whereas the spatial score reflects the organization of the local immune microenvironment. Together, these results indicate that integrating lineage and spatial immune information may improve risk stratification beyond approaches based solely on tumor-intrinsic molecular features.

These findings also have potential therapeutic implications. Classical and Hybrid tumors with immune-inflamed microenvironments may represent a subset of PDAC that is more amenable to immunotherapeutic strategies. In contrast, Basal and Null tumors, characterized by macrophage-dominated immune landscapes, may require approaches targeting the myeloid compartment, such as macrophage depletion or repolarization strategies. The SPP1–CD44 axis has previously been implicated in promoting cancer stemness within the pancreatic tumor microenvironment^38^, and our study demonstrates its selective amplification within spatially defined Hybrid_Basal niches in the context of intratumoral epithelial subtype heterogeneity. Importantly, the identification of a spatially restricted SPP1–CD44 signaling axis in Hybrid_Basal regions suggests a potential link between macrophage-mediated signaling, epithelial plasticity, and local immune remodeling. Given the established role of SPP1–CD44 signaling in tumor progression and immune suppression^37, 43^, the preferential activation of this pathway within dedifferentiated tumor niches highlights it as a candidate therapeutic target for mechanistic and therapeutic studies. In addition, the spatial immune metrics developed in this study may help refine patient selection for combination therapeutic strategies that simultaneously target tumor-intrinsic programs and the tumor microenvironment.

Several limitations of this study should be acknowledged. First, the overall cohort comprised 27 patients; however, spatial analyses were restricted to 22 patients because the remaining five did not meet the minimum threshold required for spatial metric calculation. This relatively small sample size limits statistical power for multivariable modeling, subgroup analyses, and robust assessment of prognostic biomarkers. Second, molecular subtype classification relied on cohort-derived expression thresholds, as no externally validated cutoffs currently exist for multiplex IF-based PDAC subtyping. Although the resulting subtype distribution was consistent with previously reported proportions^23^, validation in larger independent cohorts with standardized classification criteria will be necessary to confirm the robustness and generalizability of these subtype assignments. Third, CosMx spatial transcriptomic profiling was performed in only three patients, limiting transcriptomic validation to a small number of representative samples, and restricting our ability to fully capture the diversity of advanced-stage PDAC. The absence of an independent validation cohort represents an additional limitation, although publicly available multiplex spatial imaging datasets with matched clinical outcomes for advanced-stage PDAC remain scarce. Furthermore, the observational nature of this study precludes definitive conclusions regarding causal relationships between epithelial lineage state, spatial immune organization, and macrophage-mediated signaling. Finally, while our focus on advanced-stage disease enhances the clinical relevance of the study, it precludes direct comparison of immune spatial architecture between early- and late-stage PDAC.

In summary, this study provides a spatially resolved characterization of advanced PDAC, demonstrating that tumor lineage state, intratumoral epithelial heterogeneity, and spatial immune organization are closely intertwined and jointly influence clinical outcome. By integrating multiplex spatial proteomics with spatial transcriptomics, we reveal that Hybrid tumors comprise distinct epithelial and immune niches associated with differential macrophage enrichment and localized SPP1–CD44 signaling programs. Importantly, spatial immune organization provided prognostic information beyond conventional immune cell abundance metrics, and its integration with GATA6 expression achieved superior risk stratification compared with either feature alone. Together, these findings support a model in which tumor-intrinsic lineage programs and the spatial architecture of the tumor microenvironment interact to shape clinical outcome and represent complementary dimensions of PDAC biology. This lineage–immune spatial framework may inform future biomarker development and guide therapeutic strategies that simultaneously target epithelial plasticity and the immunosuppressive tumor microenvironment in advanced PDAC.

## Methods

### Study population and study design

We retrospectively collected 27 formalin-fixed, paraffin-embedded (FFPE) tissue samples from patients with locally advanced or metastatic pancreatic adenocarcinoma who were treated with standard chemotherapy at Siriraj Hospital between 2017 and 2021. Samples were collected and used with authorization from the Siriraj Institutional Review Board (SiRB), Faculty of Medicine, Siriraj Hospital, Mahidol University (COA no. 251/2565; IRB3). Overall survival (OS) was defined as the period from diagnosis to death or last follow-up. The clinical characteristics of the patients are presented in Table 1.

### Tissue preparation and staining using tissue cyclic immunofluorescence (t-CyCIF) assay

Formalin-fixed and paraffin-embedded (FFPE) tissue slides were sectioned with 5-µm thickness and prepared for t-CyCIF assay^44^. Antibody details are provided in Supplementary Table 1. Briefly, we incubated the slides in a 60°C oven for 1h, then deparaffinized them in xylene, rehydrated them through an alcohol series for 3 minutes each, and washed them again with PBS, pH 7.4. Slides were then placed in a 500 ml beaker filled with 150 ml 1xDAKO solution, pH 6.0, boiled at 90°-100°C prior to use. Antigen retrieval was performed with preheated DAKO solution for 20 min. The slides were cooled down and then blocked with 50-200 µl Odyssey blocking buffer for 1 h. After blocking, photo bleaching was performed to inactivate the autofluorescence using a solution of 4.5% H2O2 and 24 mM in the presence of white light, and finally, samples were stained and incubated with antibodies at 4°C overnight. Nuclear staining with Hoechst was followed by mounting using 10% glycerol in 1x phosphate-buffered saline (PBS; pH 7.4). Coverslips were released from slides by placing slides in (PBS; pH 7.4) for at least 15 min and the fluorophores were inactivated with 4.5% H2O2 and 24 mM NaOH under the white light every imaging cycle.

### Image acquisition

Imaging was performed using the Rarecyte HT II Finder, and the images were registered by ASHLAR software to produce a single image (ome.tiff) file and then imported into Qupath^45^ for image visualization, initial analysis, and quality control. Regions of interest (ROIs) were identified according to the instructions of a trained pathologist.

### Data integration and cell phenotyping

Registered images were loaded into QuPath v0.4.3, with two to four ROIs per sample selected to capture the largest available tumor regions per sample. Nuclear segmentation was performed on the DAPI channel using the StarDist extension (cell expansion 3 μm, threshold 0.5), and ROIs with suboptimal image quality or inconsistent marker signal were excluded prior to downstream analysis. Tissue compartments were classified as tumor or stroma based on Pan-cytokeratin staining using Gaussian mixture model with intensity thresholds reviewed across representative cases to ensure consistent classification. Of 19 panel markers, 16 markers with reliable signal intensity across the cohort were retained for cell phenotyping; three markers exhibiting high background or indistinguishable signals (CD11c, CD56, CD3d) were excluded to avoid unreliable density estimates. Retained markers included CD4, CD8, and FOXP3 (T cell subsets); CD20 (B cells); CD68 and CD163 (macrophages); PD1, PDL-1, and TIM-3 (immune checkpoint markers); CD31 (endothelial cells); SMA (stroma); PK (epithelium); GATA6 and CK5 (molecular subtype); Ki67 (proliferation); and granzyme B (cytotoxic function). Cell types were assigned using lineage marker combinations applied uniformly across all samples (Table 2). The t-CyCIF panel was developed for unautomated eight-antibody staining cycles (Supplementary Table 1). Of 27 patients, 20 had sufficient tissue for single-section staining, and 7 required distributions across two consecutive tissue sections. For cellular neighborhoods and cell–cell interaction analyses, four single-section cases (Null subtype (n=2), Classical (n=2)) were excluded to achieve balanced molecular subtype representation (n=4 per subtype), yielding a final analytical cohort of 16 patients (Supplementary Table 2). Cycle 1 staining was always performed with GATA6 and CK5. Pan-cytokeratin (PK) staining was included on both for tumor compartment classification and cross-section correspondence, with immune marker densities calculated independently per section without cross-section cell-level merging. Cell phenotyping employed a single-measurement classifier to distinguish marker-positive from marker-negative populations, with a composite classifier used for double-positive identifications. Cell densities were calculated per each ROI and normalized by area (cells/mm²). Single-cell data comprising cell type, XY coordinates, per-marker intensity, and image ID were exported as .csv files for downstream computational analysis.

### Molecular subtypes classification

Molecular subtypes were defined based on nuclear GATA6 and cytoplasmic CK5 expression in tumor cells (n=27). Pan-cytokeratin (PK) is generally used as a tumor marker. The proportion of GATA6^+^, CK5^+^, and GATA6^+^CK5^+^ colocalized cells was quantified per sample with a selected ROI. Cohort mean expression levels were used as classification thresholds (GATA6: 21.9%, CK5: 15.4%). Samples were designated Classical (GATA6-high/CK5-low), Basal (CK5-high/GATA6-low), or Null (CK5-low/GATA6-low) accordingly. Samples in which ≥5.0% of tumor cells (PK+) exhibited GATA6+CK5+ colocalization were classified as Hybrid (n=7); this threshold was set conservatively above the cohort mean coexpression rate (4.3%) to minimize misclassification from low-level non-specific colocalization. All thresholds were defined using cohort mean expression values, as externally validated cutoffs for multiplex IF-based subtyping in PDAC have not been established.

### Univariate analysis of tumor and stroma cell density

Univariable Cox proportional hazards model was performed to assess the association between cell type abundance (density/mm^2^) and overall survival in both tumor and stromal compartments separately. For each cell type, patients were dichotomized into high and low groups based on the 25th and 75th percentiles of cell abundance, with intermediate values excluded to ensure a clear contrast between groups. Hazard ratios (HR) with 95% confidence intervals and p-values were reported for each cell type. Kaplan-Meier survival curves were generated, and group differences were assessed using the log-rank test. A p-value of < 0.05 was considered statistically significant.

### Neighborhood Analysis

Cellular neighborhoods (CNs) were identified using Cytomap v1.4.21 (https://cytomap.org). Single-cell data, including X-Y coordinates and phenotype classifications, were analyzed using Raster Scan Neighborhoods with a 50 µm radius to assess local cellular composition across the cohort. Neighborhoods were clustered into eight distinct spatial microenvironments using Self-Organizing Maps (SOM). The selection of 50 µm radius was based on prior literature^46^, capturing the immediate tissue microenvironment. For cellular neighborhoods composition and cell–cell interaction analysis, we performed in 16 patients, including four cases per molecular subtype (Classical, Hybrid, Basal, Null), to avoid composition bias.

### Cellular Neighborhood Interaction Network

Cellular neighborhood (CN)–CN spatial interactions were measured by calculating mean minimum distances between all CN pairs for each molecular subtype (Classical, Hybrid, Basal, Null; n = 4 patients per subtype) using igraph. For each pair, up to 100 randomly sampled cells from one CN were used to compute Euclidean distances to all cells in another CN. Only CN pairs with at least 10 cells per CN were included, and pairs with mean distances below 50 µm were defined as CN distance. Node size was scaled to the total number of cells per CN, and the percent shared boundary as edge weight was used to describe the fraction of which CN pairs were adjacent to each other (a cutoff of 50% was used to define strongly interacting pairs). Only CN pairs present in at least three of four patients (≥75%) within a subtype were selected for data interpretation. A common layout was applied using the Fruchterman–Reingold force-directed algorithm and visualized with ggraph and ggplot2, with edge line widths continuously scaled by the shared boundary percentage and strongly interacting edges highlighted with a distinct visual encoding. Each node color represents each CN type.

### Spatial Cell-Cell Interaction Analysis

Spatial cell–cell interactions were analyzed using a neighborhood-based analytical scoring approach adapted from histoCAT ^47^ and implemented in R using the RANN package, with X/Y pixel coordinates derived from multiplex immunofluorescence imaging. For each cell-type pair per patient, spatial interaction scores were independently computed using two complementary neighborhoods definitions: a fixed-radius method (30 pixels) and a k-nearest neighbor method (k=10), following the neighborhoods framework implemented in SCIMAP^48^. Averaging scores from both methods reduced sensitivity to local cell density variation and boundary effects inherent to single-method approaches. The expected count for each ordered cell-type pair was computed. For the KNN method, a symmetric null model was applied: the expected count was computed as the product of cell type A’s global proportion, cell type B’s global proportion, and the total number of neighbor pairs. For the radius method, an asymmetric null model was applied: the expected count was computed as the total number of neighbors emitted from all cells of type A multiplied by the global proportion of cell type B, reflecting variable neighborhood sizes similar to the radius-based method.

For each ordered cell-type pair, a spatial interaction score was computed as:

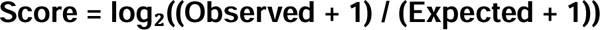

where the +1 Laplace pseudo count in both numerator and denominator ensures numerical stability for rare cell-type pairs. Pairs where both observed and expected counts which were zero were assigned as a score of zero. Positive scores indicate spatial co-localization; negative scores indicate spatial avoidance. Cell-type pairs observed in fewer than 3 patients were excluded from statistical testing. Subgroup analyses were conducted separately for Classical+Hybrid (n=8) and Null+Basal (n=8) molecular subtypes. Within each subgroup, interactions deviating significantly from spatial randomness were identified using one-sample t-tests against zero on patient-level interaction scores, with Benjamini–Hochberg FDR correction applied separately within each subtype. Pairs with FDR-adjusted p < 0.05 and an absolute mean interaction score > 0.3 were considered statistically significant. Mean interaction scores were derived from each patient, averaged across both neighborhood methods. Mean interaction score was displayed with mean ± standard error of the mean (SEM), to compare the spatial interaction patterns between molecular subgroups, though no formal between group statistical comparison was performed. Significance tiers are denoted as * FDR < 0.05, ** FDR ≤ 0.01, *** FDR ≤ 0.001.

### Spatial Score with survival Association

Building on the spatial cell-cell interaction analysis, nine candidate spatial suppression scores were defined based on positive cell type pairs demonstrating significant co-localization, using biologically relevant combinations of target cells (CD4⁺ T cells, tumor cells, B cells, endothelial cells) and suppressor cells (CD163⁺ M2 macrophages, regulatory T cells). Spatial relationships were quantified using X/Y pixel coordinates from multiplex immunofluorescence imaging. For each anchor cell (CD8⁺ or CD4⁺ T cell), the mean minimum distance to all cells of each target and suppressor phenotype within the same tissue section was calculated. A spatial suppression score was defined for each three-cell-type combination as:

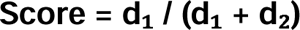

where d₁ is the mean minimum distance from the anchor cell to the nearest another target cell, and d₂ is the mean minimum distance from the anchor cell to the nearest suppressor cell. Higher scores indicate anchor cells positioned relatively closer to immunosuppressive cells and farther from target cells, reflecting a spatially unfavorable immune configuration. For each of the nine candidate scores, univariate Cox proportional hazards regression was performed on each continuous score to estimate hazard ratio, 95% CI, and concordance index. Patients were additionally dichotomized at the median for Kaplan–Meier visualization and log-rank testing. Both analyses were conducted for all nine candidate scores without multiple testing correction. The broader cohort of 27 patients with available coordinate data was used and calculated; however, of whom 22 met the minimum cell threshold (≥5 cells per type) and were included in spatial score and survival analysis.

### Spatial Transcriptomic Analysis

Formalin-fixed paraffin-embedded (FFPE) tissue sections (5 µm) were processed using the CosMx Spatial Molecular Imager (NanoString) with a 1,000-gene targeted RNA panel, following the manufacturer’s protocol^49^. Sections were imaged, segmented at single-cell resolution, and transcripts were assigned to individual cells based on in situ hybridization signals. Epithelial malignant cells were selected using cell-type annotations and analyzed in Seurat following normalization, variable feature selection, and supervised clustering. Dimensionality reduction was performed using principal component analysis and UMAP. Classical and Basal transcriptional scores were computed per malignant cell using AddModuleScore (Seurat; ctrl = 20, seed = 42) with curated Classical and Basal signature gene lists. A Delta score was defined per cell as Classical score − Basal score. FOV-level scores were derived by averaging per cell scores across all cells within each FOV and only FOVs with ≥10 malignant cells were collected. FOVs were assigned to four transcriptional lineage groups using a two-step classification rule: FOVs from molecularly classified Classical or Basal tumors were assigned accordingly based on mIF subtype annotation, while FOVs from Hybrid tumors were further subdivided by Delta score directionality (Hybrid_Classical: Delta ≥ 0.1; Hybrid_Basal: Delta < 0.1). Supervised clustering of Malignant cells from four lineage groups was performed using top marker genes and DEGs hybrid lineage groups. Scaled expression of top lineage markers was overlaid on the supervised UMAP using per-gene diverging color scales anchored at the 99th percentile of absolute scaled expression values. Macrophages were identified from CosMx cell type annotations and subset from the full cellular dataset. Unsupervised clustering was performed using a shared nearest-neighbor graph (FindNeighbors) and cluster marker genes were identified using FindAllMarkers (Wilcoxon rank-sum test; min.pct = 0.1, log₂FC threshold = 0.25). Top markers per cluster were selected by average log₂FC among genes with BH-adjusted p < 0.05 and used to assign functional macrophage states.

### Gene Set Variation Analysis

Pseudobulk expression profiles were computed for Hybrid_Classical and Hybrid_Basal groups using AverageExpression on log-normalized counts. GSVA was performed using the gsvaParam method against MSigDB Hallmark gene sets (Homo sapiens)^50^. The top 10 pathways were selected by variance between them.

### Ligand–Receptor Interaction Analysis

CellChat was used to infer intercellular ligand–receptor interactions^51^, applied independently to each FOV lineage group using the secreted signaling, cell–cell contact, and ECM-receptor databases. Communication probabilities were computed using the default mass action model. Cross-group comparisons of SPP1 pathway interaction probabilities were assessed by Wilcoxon rank-sum test with Benjamini–Hochberg correction.

### Survival Analysis

Survival analyses were performed using Cox proportional hazards regression to evaluate the association of spatial, immunological, and clinicopathological variables with overall survival. The spatial score was defined as the primary spatial variable of interest. Additional continuous variables included GATA6 expression, CK5 expression, serum CA19-9, CD8⁺Ki67⁺ proliferating CD8 T cell density, and M1 macrophage density. All continuous variables were standardized to unit variance (z-scored) prior to model entry. Clinicopathological covariates included age at diagnosis (≤60 vs. >60 years), Gender (female as reference), pathological stage (Stage III vs. IV), and molecular subtype (Classical/Hybrid [Favorable, reference] vs. Null/Basal [Unfavorable]). Univariate Cox models were applied to all variables, after which three variables were simultaneously entered into a multivariable model to avoid overfitting. Only patients with complete data (n=22) were included. Results from both analyses were displayed as a single forest plot showing log-scaled HRs with 95% CIs, with p-values < 0.05, ≤ 0.01, and ≤ 0.001.

### Time-Dependent ROC Analysis

Time-dependent receiver operating characteristic (ROC) curves were estimated using the timeROC R package with inverse probability of censoring weighting (IPCW), with the censoring distribution estimated non-parametrically via the Kaplan–Meier method. Eight markers were evaluated: the spatial score, GATA6 expression, CK5 expression, CA19-9, CD8⁺Ki67⁺ cell density, M1 macrophage cell density, and two composite scores. Composite scores were derived as linear predictors from univariate Cox proportional hazards models combining the spatial score with either GATA6 or CK5 expressions, with both variables standardized prior to model fitting and Efron’s method was applied for tied event times. Summary AUC values were computed as weighted averages of time-specific estimates across the follow-up period, with weights proportional to increments of the Kaplan–Meier-derived cumulative incidence function. Uncertainty was quantified by 1,000 bootstrap resamples and Cox models were refitted on each resample. 95% confidence intervals were derived using percentile method. A prognostic score was constructed using markers selected by iAUC performance, dichotomized at the median combined score.

### Statistics and Reproducibility

Since this study is a retrospective cohort study, the sample size was not predetermined. All analyses were performed in R (v4.4.3). R packages used included Seurat, harmony, RANN, InSituType, CellChat, GSVA, msigdbr, survival, timeROC, survminer, ggplot2, ComplexHeatmap, patchwork, ggsignif, forestplot, dplyr, tidyr, reshape2, rstatix and igraph.

## Data availability

The data generated during this study are included in this published article and provided as Supplementary Information.

## Code availability

The underlying code for this study is available in GitHub (PDAC-data-analysis) and is publicly available as of the date of publication.

## Acknowledgements

This research was supported by the National Research Council of Thailand (NRCT) (2022) and the Siriraj Research Development Fund, Faculty of Medicine Siriraj Hospital, Mahidol University (Grant Nos. R016837002 and R016733025). S.S. was supported by the Research Excellence Development (RED) Program, Faculty of Medicine Siriraj Hospital, Mahidol University.

## Author contributions

S.S. conceived and supervised the study. H.M.O. and S.S. designed the experiments. H.M.O. performed the experiments and data analysis. H.M.O. and S.S. did data interpretation. T.A. did ROI selection, and N.A. and A.P. performed sample recruitment and processing. K.K. assist in providing clinical data and interpretation. U.D. did the CosMx experiment and S.P. assist in CosMx data interpretation. H.M.O. curated the dataset and prepared the figures. H.M.O. and S.S. wrote the manuscript. All authors reviewed and approved the final manuscript.

## Competing interests statement

The authors declare no competing interests.

## Notes

### Competing Interest Statement

The authors have declared no competing interest.

### Summary of Updates

Author name corrected; Fig. 2.d corrected.

