## Supplementary Figures for "Spatial immune architecture and tumor lineage programs jointly shape clinical outcomes in advanced pancreatic ductal adenocarcinoma"

### Supplementary Information

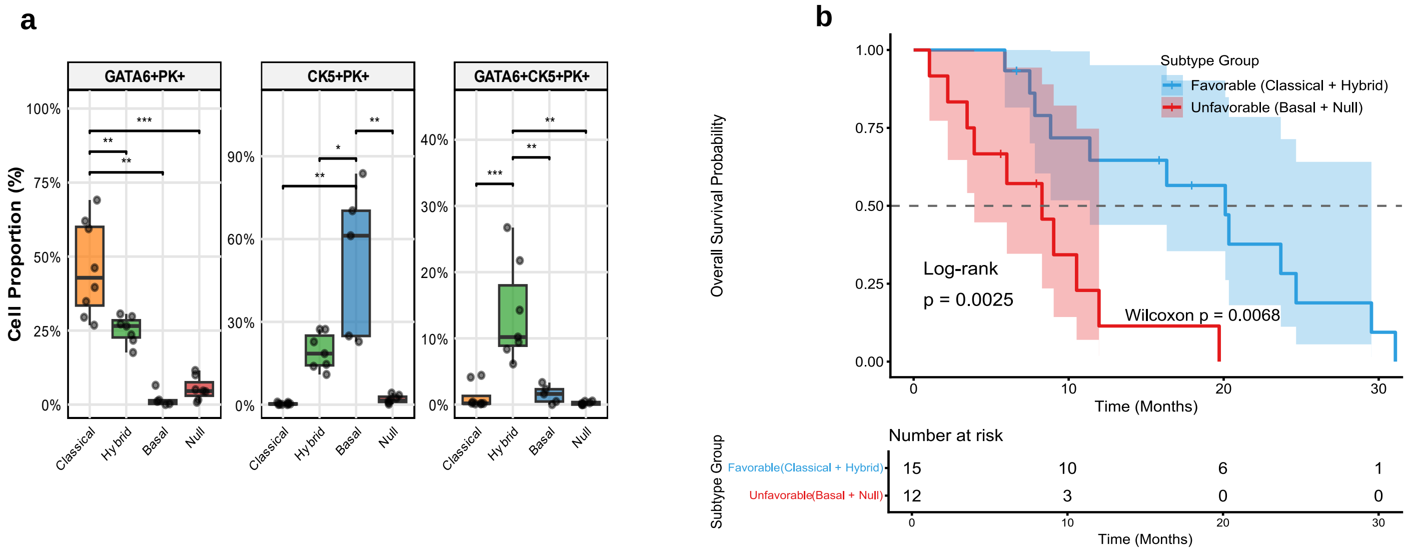


**Supplementary Figure 1: Molecular marker expression in distinct molecular subtypes. a** Boxplots showing the proportion of GATA6+PK+, CK5+PK+, and GATA6+CK5+PK+ cells across molecular subtypes. Boxes represent the interquartile range (IQR) with the median line; dots indicate individual patient values. **b** Kaplan–Meier survival analysis comparing Favorable and Unfavorable patient groups, with significance assessed by log-rank and Wilcoxon rank-sum tests. $*p < 0.05$, $**p < 0.01$, $***p < 0.001$.

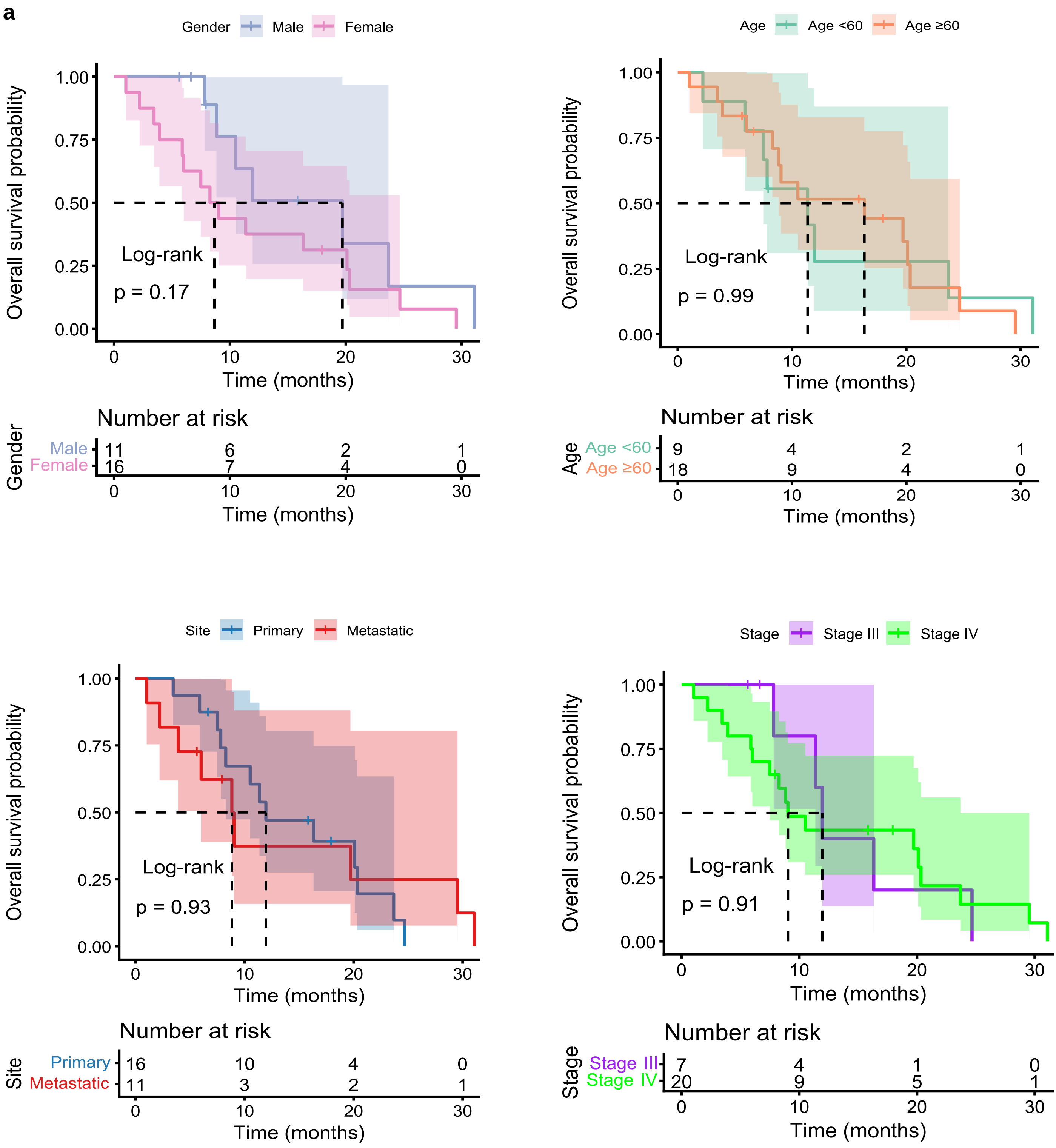
a
Gender Male Female
Overall survival probability
1.00
0.75
0.50
0.25
0.00
Log-rank
p = 0.17
0 10 20 30
Time (months)
Number at risk
Gender
Male 11 6 2 1
Female 16 7 4 0
0 10 20 30
Time (months)
Age Age <60 Age ≥60
Overall survival probability
Log-rank
p = 0.99
Time (months)
Number at risk
Age
Age <60 9 4 2 1
Age ≥60 18 9 4 0
Time (months)
Site Primary Metastatic
Overall survival probability
Log-rank
p = 0.93
Time (months)
Number at risk
Site
Primary 16 10 4 0
Metastatic 11 3 2 1
Time (months)
Stage Stage III Stage IV
Overall survival probability
Log-rank
p = 0.91
Time (months)
Number at risk
Stage
Stage III 7 4 1 0
Stage IV 20 9 5 1
Time (months)

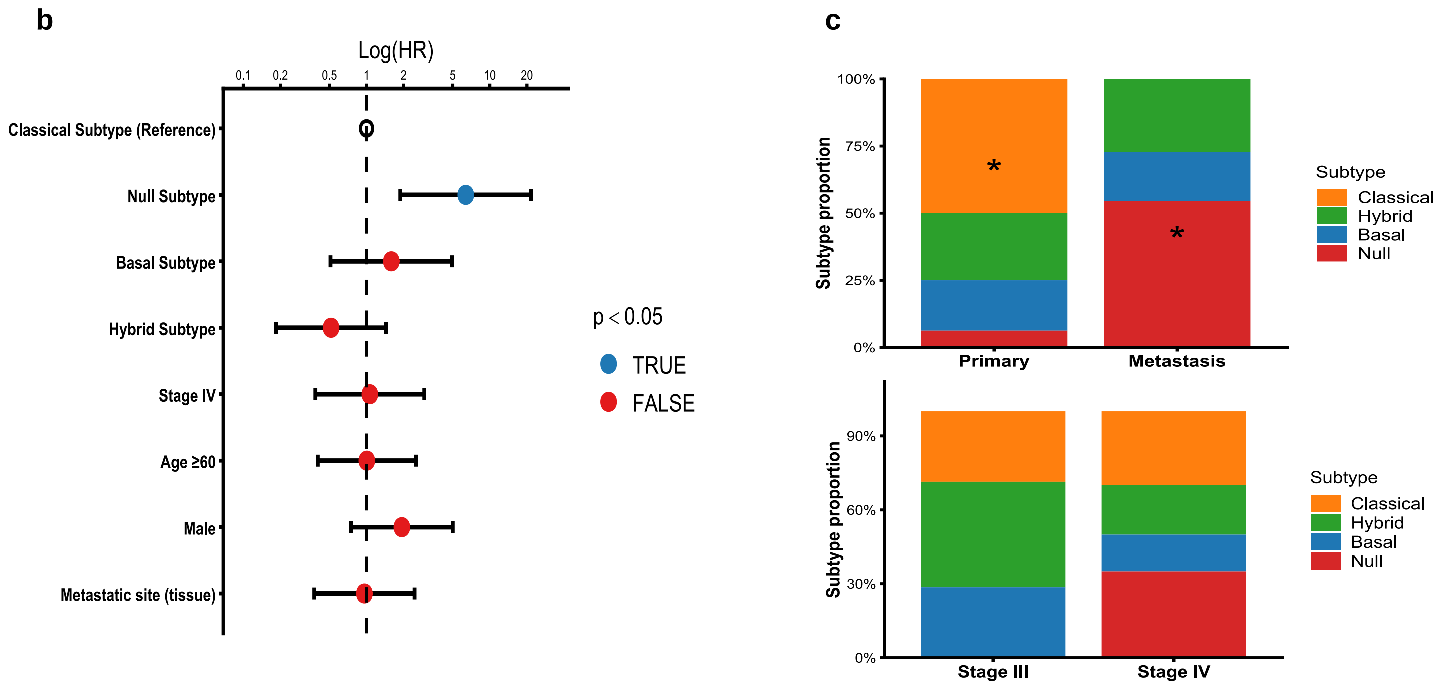


**Supplementary Figure 2: Prognosis association of molecular subtypes and clinical variables in cohort.** **a** Association of clinical variables with patient outcomes: gender, age at diagnosis, site of tissue and stage at diagnosis. **b** Forest plot showing hazard ratios (HR) and 95% confidence intervals overall survival across clinical and molecular subgroups on log scale. Blue points indicate $p < 0.05$; red points indicate $p \geq 0.05$. **c** Stack bar plot comparing different sites and stages of PDAC among subtypes. Survival analysis was conducted using Kaplan-Meier method with log-rank test for significance. *P* values < 0.05 were considered statistically significant**.**

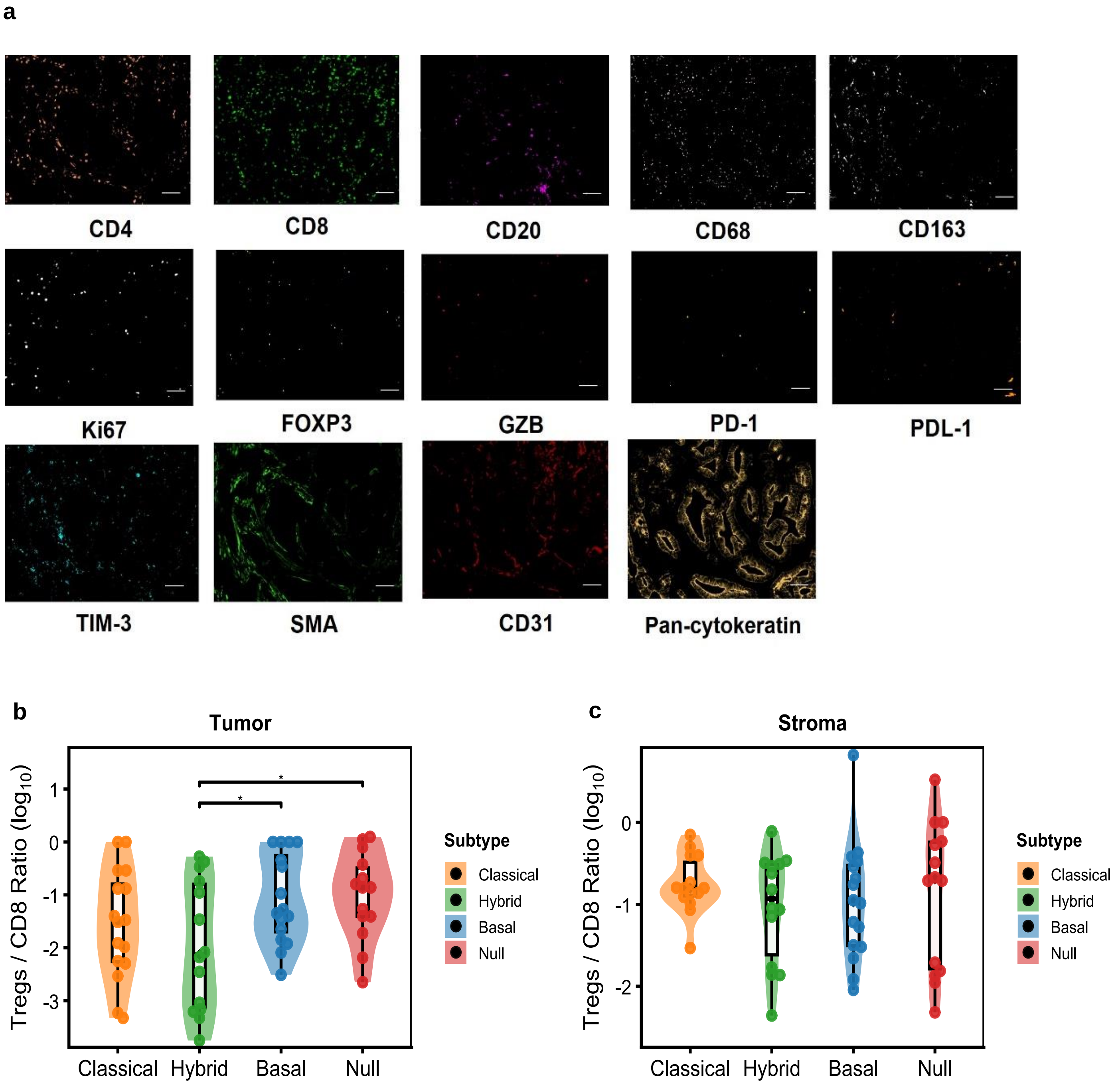


**Supplementary Figure 3: Tumor microenvironment characterization across molecular subtypes. a** Representative multiplex immunofluorescence marker panels demonstrating expression patterns (scale bar-50µm). **b-c** Tregs/CD8 ratios across molecular subtypes in Tumor and Stroma compartments. Statistical significance was assessed by Kruskal-Wallis's test with post-hoc pairwise comparisons. $*p < 0.05$, $**p \leq 0.01$, $***p \leq 0.001$.

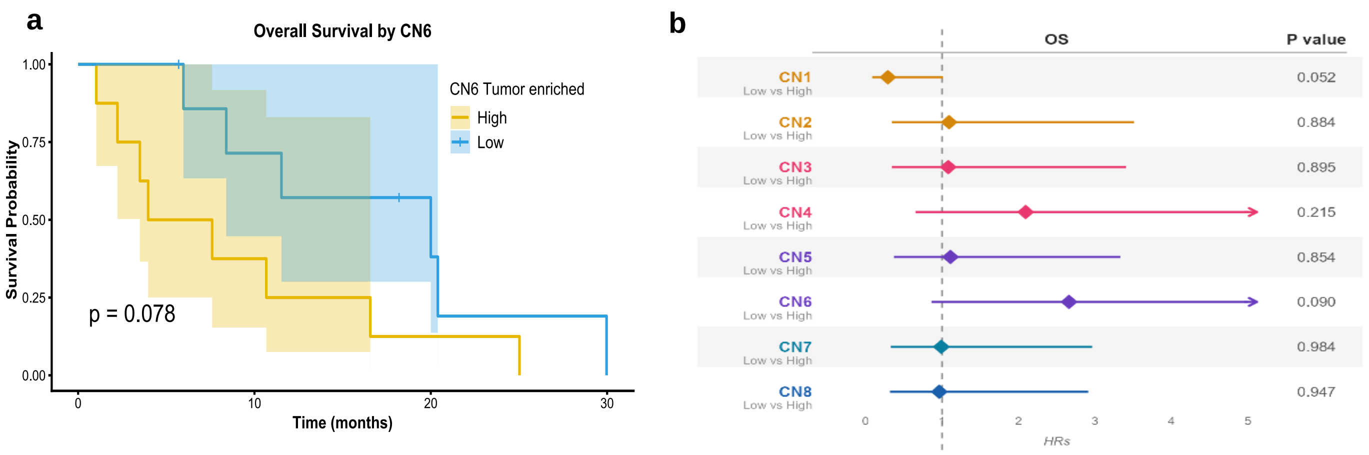


**Supplementary Figure 4: Cellular neighborhoods and survival associations across molecular subtypes.** **a** Kaplan–Meier overall survival curves stratified Tumor-enriched neighborhoods (CN6) frequencies, dichotomised at the median value. Survival analysis was conducted using log-rank test for significance. **b** Forest plot showing hazard ratios (HR) and 95% confidence intervals for overall survival of all cellular neighborhoods. *P* values < 0.05 were considered statistically significant**.** Shaded areas in KM curve represent 95% confidence intervals.

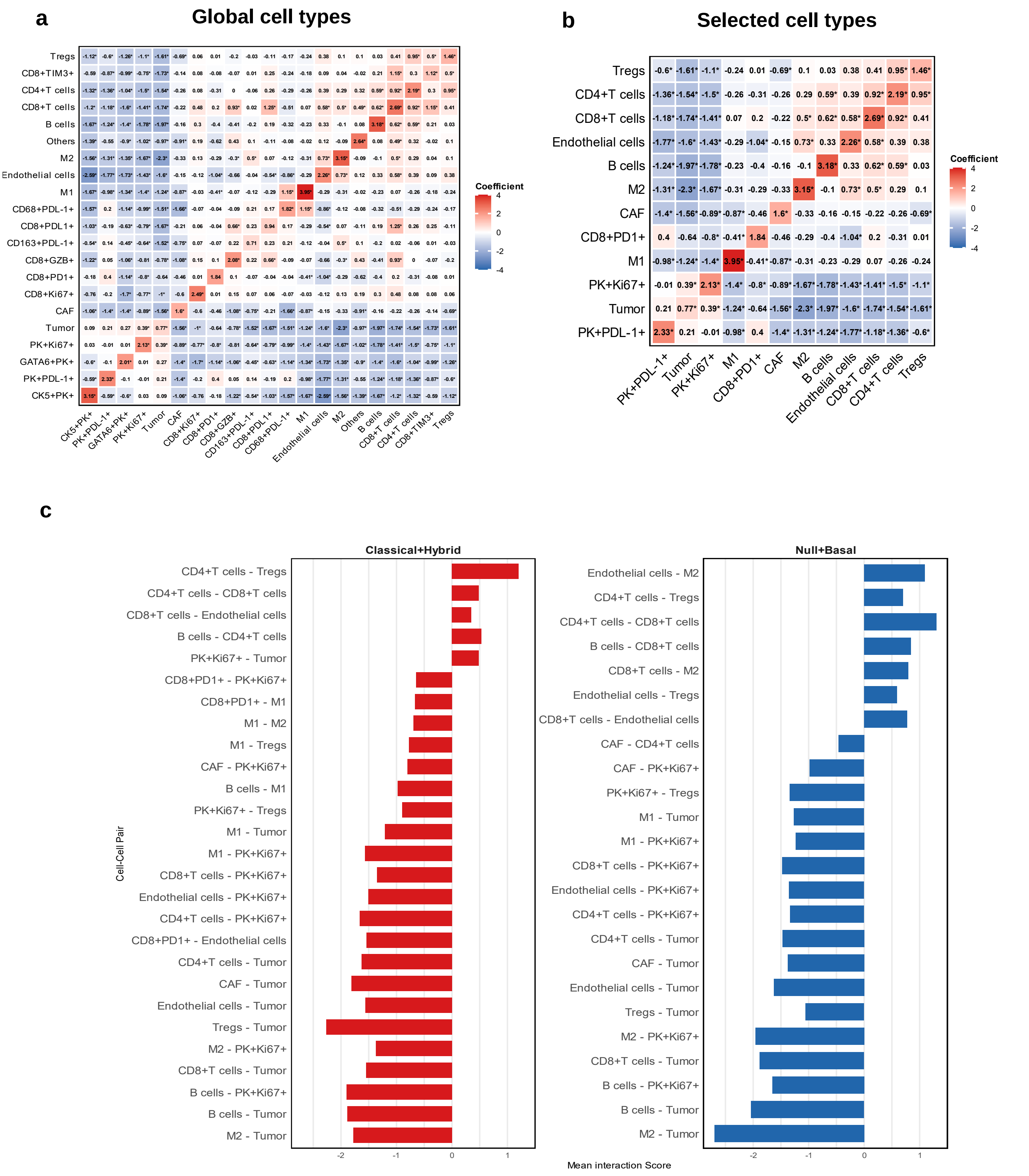


**Supplementary Figure 5: Cell to cell interaction analysis across molecular subtypes. (a-b)** Heatmap of pairwise spatial interaction scores in global cell populations and selected cell populations across all samples (n=16), with cell types ordered by hierarchical clustering using Euclidean distance. **c** Mean interaction scores (±SEM) for statistically significant cell-cell pairs in Classical+Hybrid (red) and Null+Basal (blue) subgroups, displayed side by side for visual comparison. For all panels, significance was determined using one-sample t-tests against zero with Benjamini–Hochberg FDR correction within each subgroup; pairs with FDR < 0.05 and absolute mean interaction score > 0.3 were considered as statistically significant. Significance tiers: * FDR < 0.05, ** FDR ≤ 0.01, *** FDR ≤ 0.001.

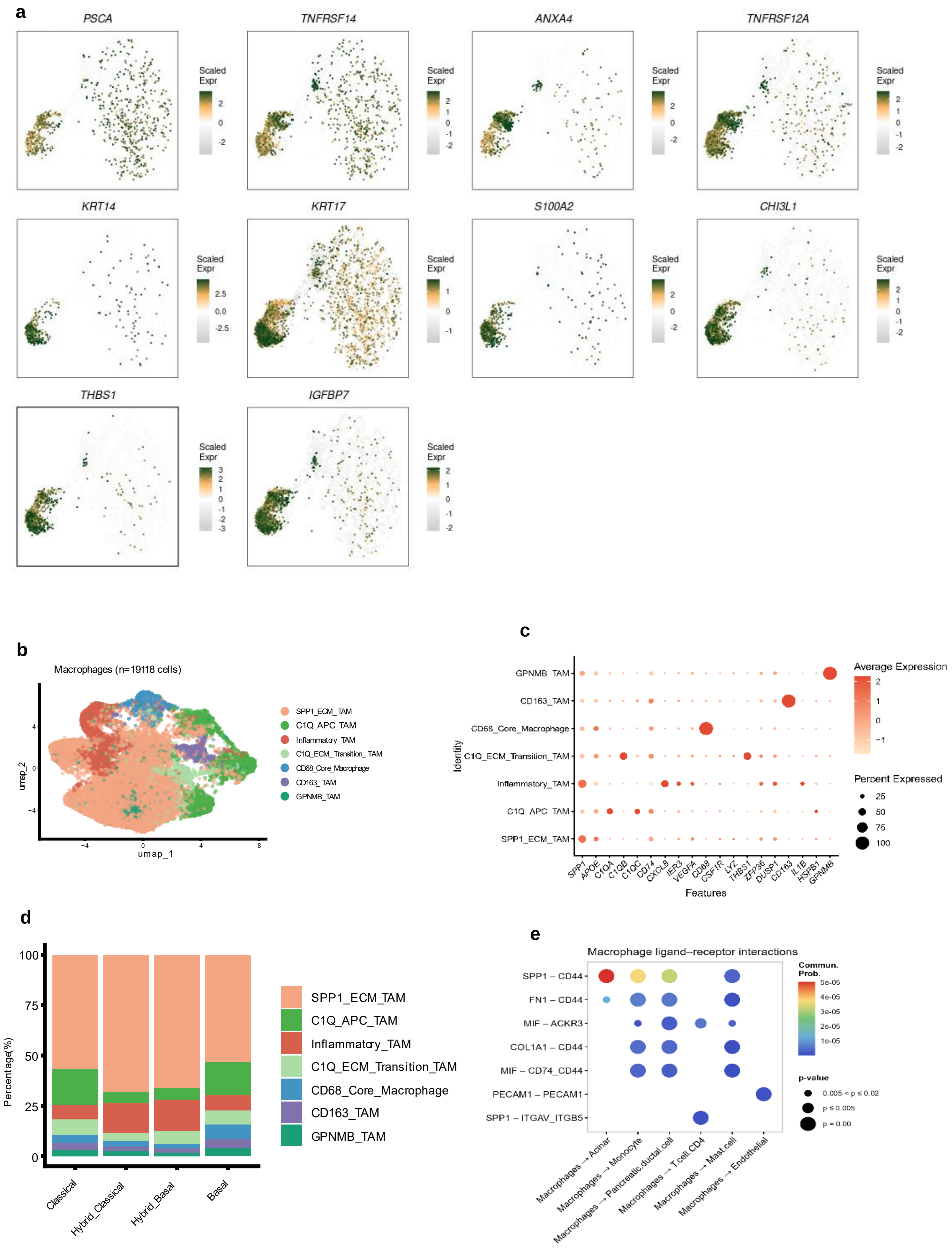

a
PSCA
TNFRSF14
ANXA4
TNFRSF12A
KRT14
KRT17
S100A2
CHI3L1
THBS1
IGFBP7
Scaled Expr
b
Macrophages (n=19118 cells)
umap_1
umap_2
SPP1_ECM_TAM
C1Q_APC_TAM
Inflammatory_TAM
C1Q_ECM_Transition_TAM
CD68_Core_Macrophage
CD163_TAM
GPNMB_TAM
c
GPNMB_TAM
CD163_TAM
CD68_Core_Macrophage
C1Q_ECM_Transition_TAM
Inflammatory_TAM
C1Q_APC_TAM
SPP1_ECM_TAM
Identity
SPP1 APOE C1QA C1QB C1QC CD74 CXCL8 IER3 VEGFA CD68 CSF1R LYZ THBS1 ZFP36 DUSP1 CD163 IL1B HSPB1 GPNMB
Features
Average Expression
Percent Expressed
25
50
75
100
d
Percentage(%)
Classical
Hybrid_Classical
Hybrid_Basal
Basal
SPP1_ECM_TAM
C1Q_APC_TAM
Inflammatory_TAM
C1Q_ECM_Transition_TAM
CD68_Core_Macrophage
CD163_TAM
GPNMB_TAM
e
Macrophage ligand−receptor interactions
SPP1 − CD44
FN1 − CD44
MIF − ACKR3
COL1A1 − CD44
MIF − CD74_CD44
PECAM1 − PECAM1
SPP1 − ITGAV_ITGB5
Macrophages → Acinar
Macrophages → Monocyte
Macrophages → Pancreatic.ductal.cell
Macrophages → T.cell.CD4
Macrophages → Mast.cell
Macrophages → Endothelial
Commun. Prob.
5e-05
4e-05
3e-05
2e-05
1e-05
p-value
0.005 < p ≤ 0.02
p ≤ 0.005
p = 0.00

**Supplementary Figure 6: Intratumoral spatial heterogeneity within Hybrid subtype. a** Feature plots showing scaled expression of representative marker genes on the supervised UMAP. Color scale represents scaled expression (z-scored) per gene independently; blue, low; white, average; orange, moderate-high; dark green, highest expression. **b** UMAP visualization of macrophage populations (n = 19,118 cells) from CosMx profiling, annotated by marker gene expression. **c** Dot plot showing normalized expression of marker genes across seven transcriptionally distinct macrophage subtypes. Dot size reflects the percentage of cells expressing each gene; color intensity reflects mean scaled expression. **d** Stacked bar plot showing proportional distribution of macrophage subtypes across Classical, Hybrid_Classical, Hybrid_Basal and Basal tumors, demonstrating enrichment of SPP1/ECM and inflammatory TAM populations in Hybrid tumors**. e** Bubble plot of macrophage ligand–receptor interactions with indicated cell types. Bubble color reflects communication probability (blue to red: low to high); bubble size reflects statistical significance (larger = lower p-value).
